# Elicitation of stem-directed antibodies in rhesus macaques by a conventional hemagglutinin immunogen

**DOI:** 10.64898/2026.07.16.738984

**Authors:** Shuqin Gu, Joel Finney, Kan Luo, Sarah M. Valencia, Dieter Mielke, Tarra A. Von Holle, Dawn J. Marshall, Robert Parks, Laura L. Sutherland, Richard M. Scearce, Kevin Wiehe, Sampa Santra, Summer Harris, Jacquelyn DuVall, Chelsea D. Landon, M. Ariel Spurrier, Nicholas S. Heaton, Hadi M. Yassine, Barney S. Graham, Thomas B. Kepler, Hua-Xin Liao, Aaron G. Schmidt, Guido Ferrari, Barton F. Haynes, Stephen C. Harrison, M. Anthony Moody

## Abstract

Because they can bind many strains of influenza, antibodies targeting the hemagglutinin (HA) stem have been attractive targets for vaccine development. Many monoclonal antibodies (mAbs) directed at the HA stem have been isolated from humans, and these mAbs have mediated broad protection in animal models. We describe here HA stem-directed mAbs isolated from rhesus macaques immunized with an “ordinary” H1 HA trimer. All immunized rhesus macaques developed high serum titers with broad reactivity to diverse H1N1 and H5N1 viruses, and 7 isolated mAbs strongly blocked canonical stem antibody CR6261 binding to H1. MAb DH726.1 robustly protected mice from lethal challenge with H1N1 and H5N1 viruses, and cryo-EM showed the binding footprint overlapped that of some human mAbs. These findings suggest that vaccination with the standard, trimeric HA immunogens may be sufficient to elicit stem antibodies at titers adequate to protect against zoonotic H5N1 influenza.

**In Brief:** Efforts to achieve broad influenza protection have largely emphasized increasingly sophisticated immunogen designs to redirect antibody responses toward conserved epitopes. Here, we show that a single, conventional HA immunogen readily elicits antibodies targeting the conserved HA stem, suggesting that routine influenza vaccination may provide broader protection than previously appreciated.

**Highlights:**

1. Monovalent HA immunization induces strong, stem-directed immune responses in rhesus macaques
2. Epitope on the HA stem confirmed by Cryo-EM
3. Fc-mediated activation of immune effector cell by rhesus-derived stem mAb
4. HA stem-directed antibody protects mice against H1N1 and H5N1 viruses

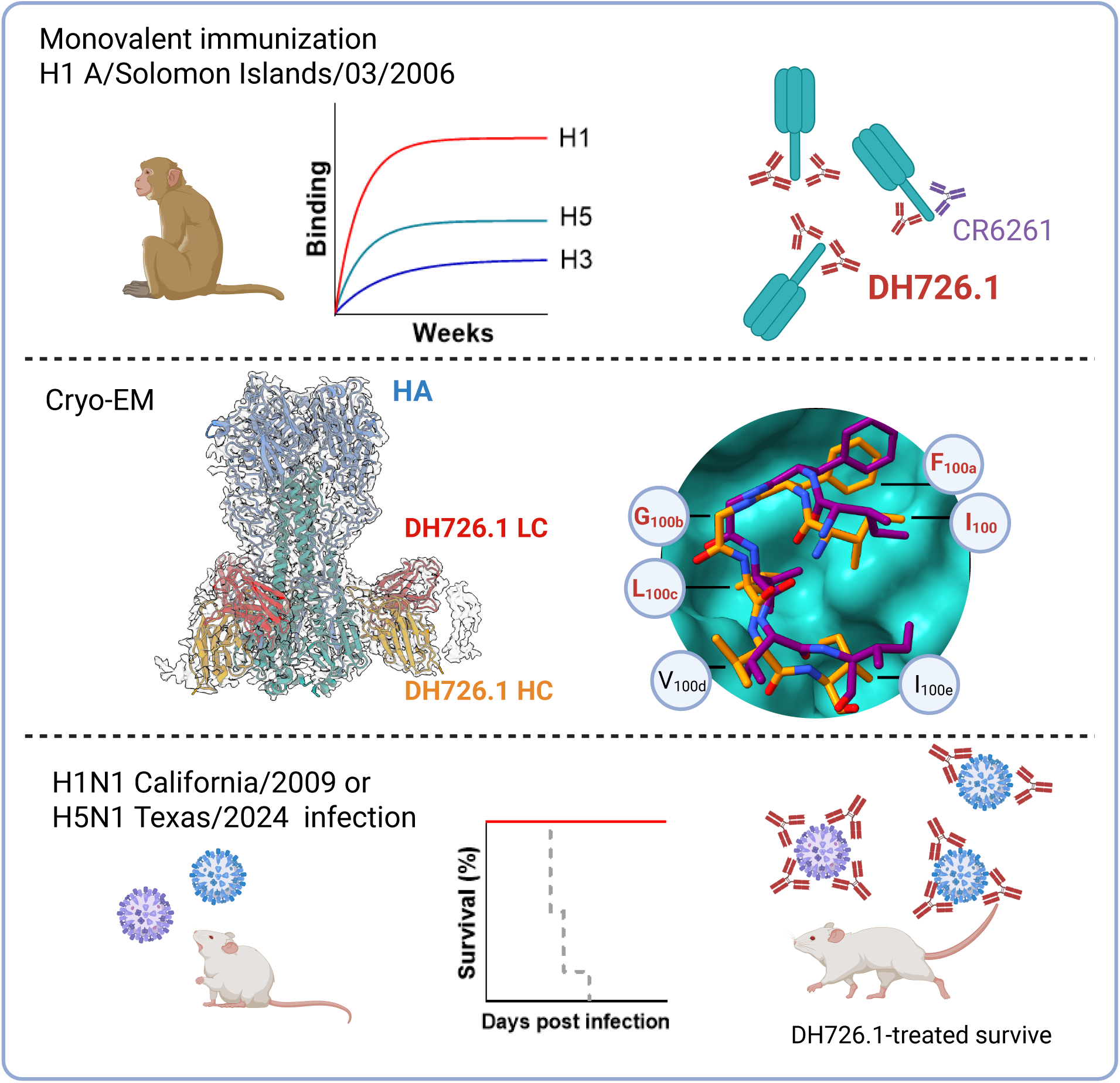

## Introduction

Influenza is a persistent health threat requiring on-going surveillance of circulating strains (*1, 2*). Vaccination lowers disease severity, slows transmission, and reduces morbidity and mortality (*3–5*), but current influenza vaccines have variable efficacy in humans, ranging from 19% to 60% (*6*). Sites on the major surface glycoprotein hemagglutinin (HA) are potential targets for broadly protective antibodies, including a conserved hydrophobic region along the HA stem (*7*) and the sialic acid binding site on the HA head (*8*). Immunogens designed to preferentially elicit antibodies to these sites are being tested as “universal” influenza vaccine candidates. Antibodies that inhibit sialic acid receptor engagement are usually subtype specific (*9, 10*); those with heterosubtypic breadth generally target epitopes conserved over a somewhat broader footprint, including those on the membrane-proximal HA stem, and there have been efforts to develop influenza vaccines that can elicit stem-directed responses (*7, 11–13*).

Testing of vaccine candidates often uses animal models such as mice or ferrets (*14*). While both mice and ferrets can develop symptomatic influenza infection, their responses may not directly mimic those of humans (*15*). In contrast, rhesus macaques are more closely related to humans (*16*), and can mimic human immune responses, as shown for Zika virus (*17, 18*), cytomegalovirus (*19*), and HIV-1 (*20, 21*). We tested whether rhesus macaques immunized with recombinant influenza HA could develop HA stem-directed antibodies like those observed in humans.

## Results

### Influenza H1 immunization of rhesus macaques elicits plasma antibodies that bind multiple influenza strains

Six rhesus macaques were immunized five times with recombinant influenza HA H1 A/Solomon Islands/03/2006 (Figure 1A). All animals developed antibodies that bound the immunizing HA (Figure 1B), the closely related H1 A/Brisbane/59/2007 (Figure 1C), and pandemic H1 A/California/04/2009 (Figure 1D), though the latter was at lower levels. We observed weaker binding to HAs from A/Wisconsin/67/2005 (H3N2) (Figure 1E) and group 1 cross-reactivity to A/Vietnam/1203/2004 (H5N1) and A/Indonesia/05/2005 (H5N1) (Figure 1F and G, respectively). Anti-HA binding was elicited by the first immunization and increased with boosting, plateauing after the first boost.

**Figure 1.**
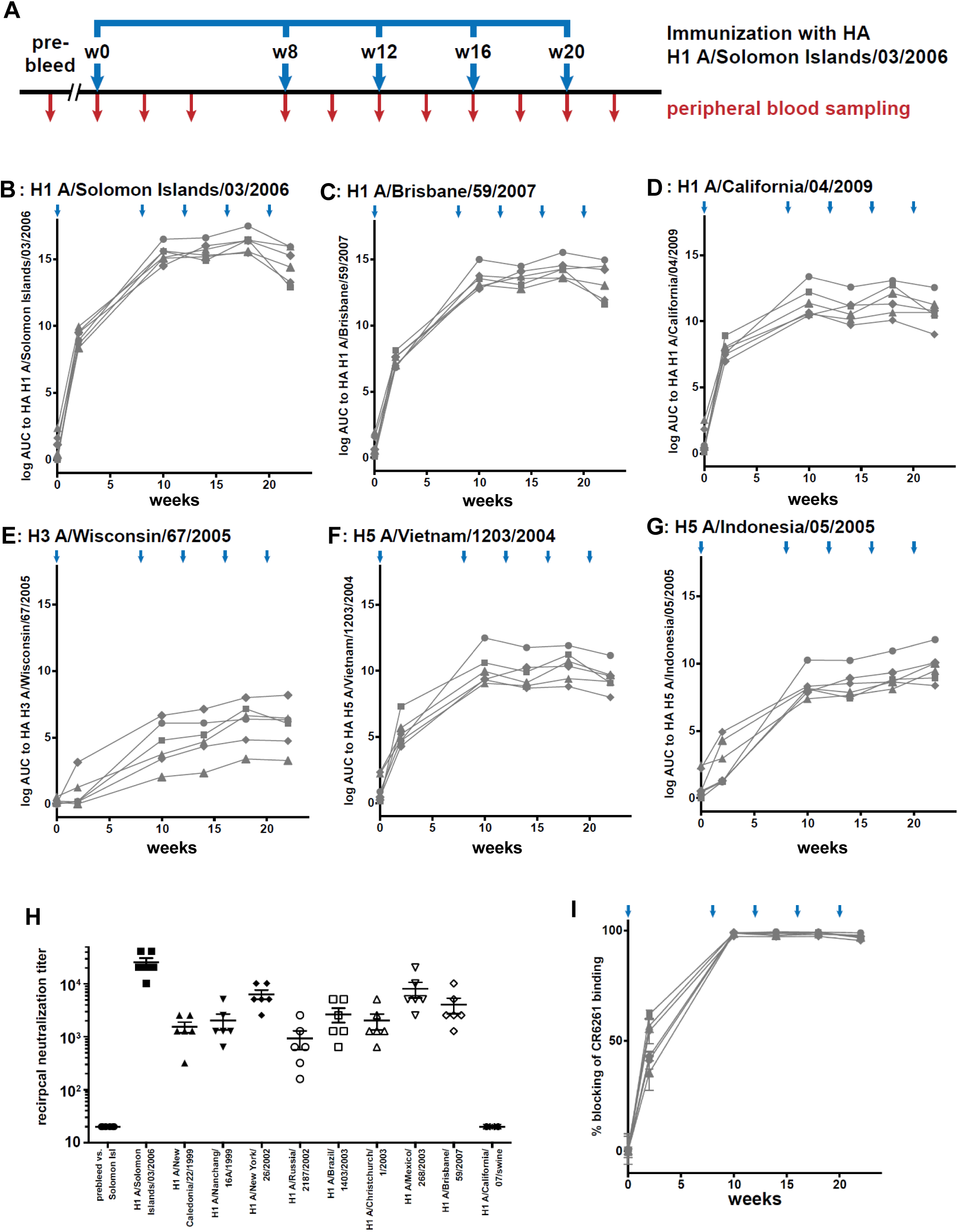
Plasma/sera antibody binding, neutralization, and blocking assays. (A) Immunization and bleed schedule aligned to the binding curves. Binding levels plotted at log area under the curve (AUC) for six different HA proteins (B-G). (H) Neutralization. Pre-immune sera were tested against H1 A/Solomon Islands/03/2006 and sera after the final immunization were tested against a panel of H1 strains. There was no activity against a strain related to the 2009 pandemic. (I) Plasma antibody blocked binding of stem-directed mAb CR6261 to H1 A/Solomon Islands/03/2006.

Plasma from rhesus macaques lacked neutralizing activity before immunization but had high titers against A/Solomon Islands/03/2006 (H1N1) after the final immunization (Figure 1H). Despite binding to the HA of the 2009 H1N1 pandemic strain, we found no neutralization of A/California/07/2009 (H1N1), suggesting that the cross-reactive antibodies detected by ELISA could not bind intact virions. This inference is consistent with work showing that antibodies directed against the HA stem can bind multiple strains of influenza but neutralize weakly (*22, 23*). Thus, we tested whether immune plasma could block binding of the stem-directed mAb CR6261. All macaques developed high levels of CR6261-blocking antibodies after the second immunization (Figure 1I). These results indicated that the vaccine regimen we used had elicited antibodies to the HA stem.

### Memory B cell receptor repertoire analysis

We isolated HA-specific B cells by modifying a technique we used to isolate HIV-1-specific B cells (*24, 25*), using fluorochrome-labelled HA reagents as “baits” to bind HA-specific surface B-cell receptors (BCRs). Our sort was designed to isolate stem-directed cross-reactive BCRs, and we used antibody-coated beads to test our four baits for specificity. None of our conjugates bound to control mAb P3X63, while 3 of 4 bound to stem-directed antibody CR6261 (Figure S1A). H1 A/Solomon Islands/03/2006 bound to receptor binding site mAb CH65 (*8*), while 3 of 4 bound to mAb DH253 (*26*). We paired H1 A/California/04/2009 and H5 A/Vietnam/1203/2004 to preferentially select B cells expressing stem-directed antibodies, and isolated HA-specific memory B cells from week 22 PBMCs by flow cytometry (Figure 2A).

**Figure 2.**
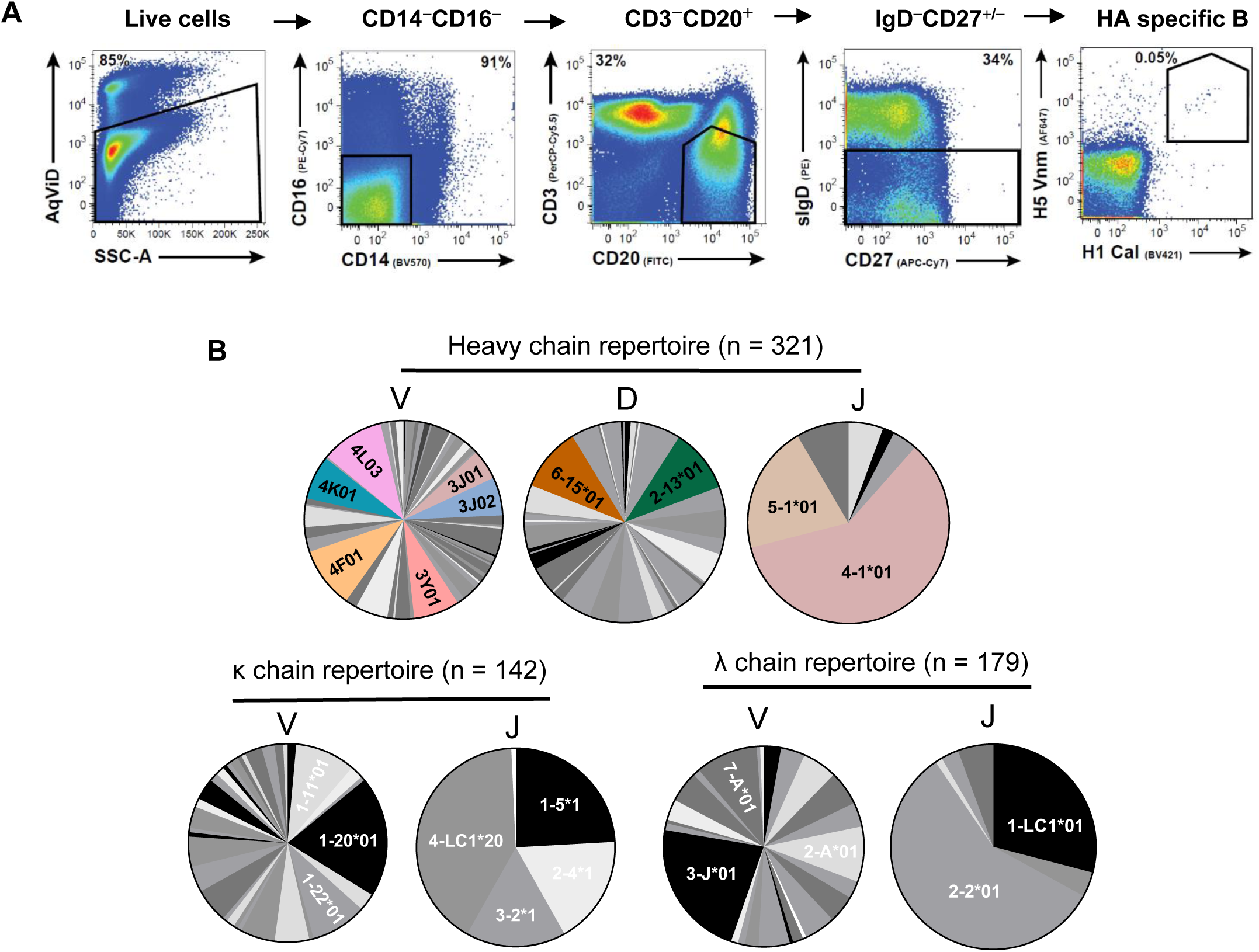
Characteristics of HA-specific B cells. (A) Flow cytometric gating strategy for isolation of antigen-specific B cells. Geometric gating for lymphocytes and singlets were not shown. (B) Ig V_H_ and V_L_ gene family repertoire of H1^+^ and H5^+^ memory B cells isolated 22 weeks post immunization. The repertoire is shown as a pie chart.

We analyzed the genes from the isolated HA-reactive B cells, and found that six heavy chain V genes *(IGHV3J01*, *IGHV3J02*, *IGHV3Y01*, *IGHV4F01*, *IGHV4K01*, *IGHV4L03)* were used by >45% of recovered B cells, two diversity segments *(IGHD2-13*01, IGHD6-15*01)* were used by >20%, and two heavy-chain joining segments *(IGHJ4-1*01, IGHJ5-1*01)* were used by most recovered B cells (Figure 2B top panel). About 20% of κ chain sequences used *IGKV1-20*01*, and more than 40% used *IGKJ4-LC1*20*; >20% of λ chains used *IGLV3-J*01*, and >50% used *IGLJ2-2*01* (Figure 2B bottom panel). The average frequency of somatic mutations was higher in heavy chains than in κ chains, but λ chains had a larger range of mutation frequencies (Figure S1B).

### Characterization of isolated stem-directed mAbs

We expressed mAbs from 20 isolated memory B cells, of which nine bound to H1 A/Solomon Islands/03/2006 (Table 1); 2/9 (22%) were clonally related. We found that 9 mAbs bound well to H1 and H5 HAs, but not H3s (Figure 3A); 7/9 (78%) mAbs blocked binding of mAb CR6261 to H1 A/Solomon Islands/03/2006 (Figure 3B). The two non-CR6261-blocking mAbs showed stronger HAI activity than the blocking mAbs, of which three exhibited low-level HAI activity against A/Wisconsin/67/2005 (H3N2) while the other four had no HAI activity (Figure 3C).

**Figure 3.**
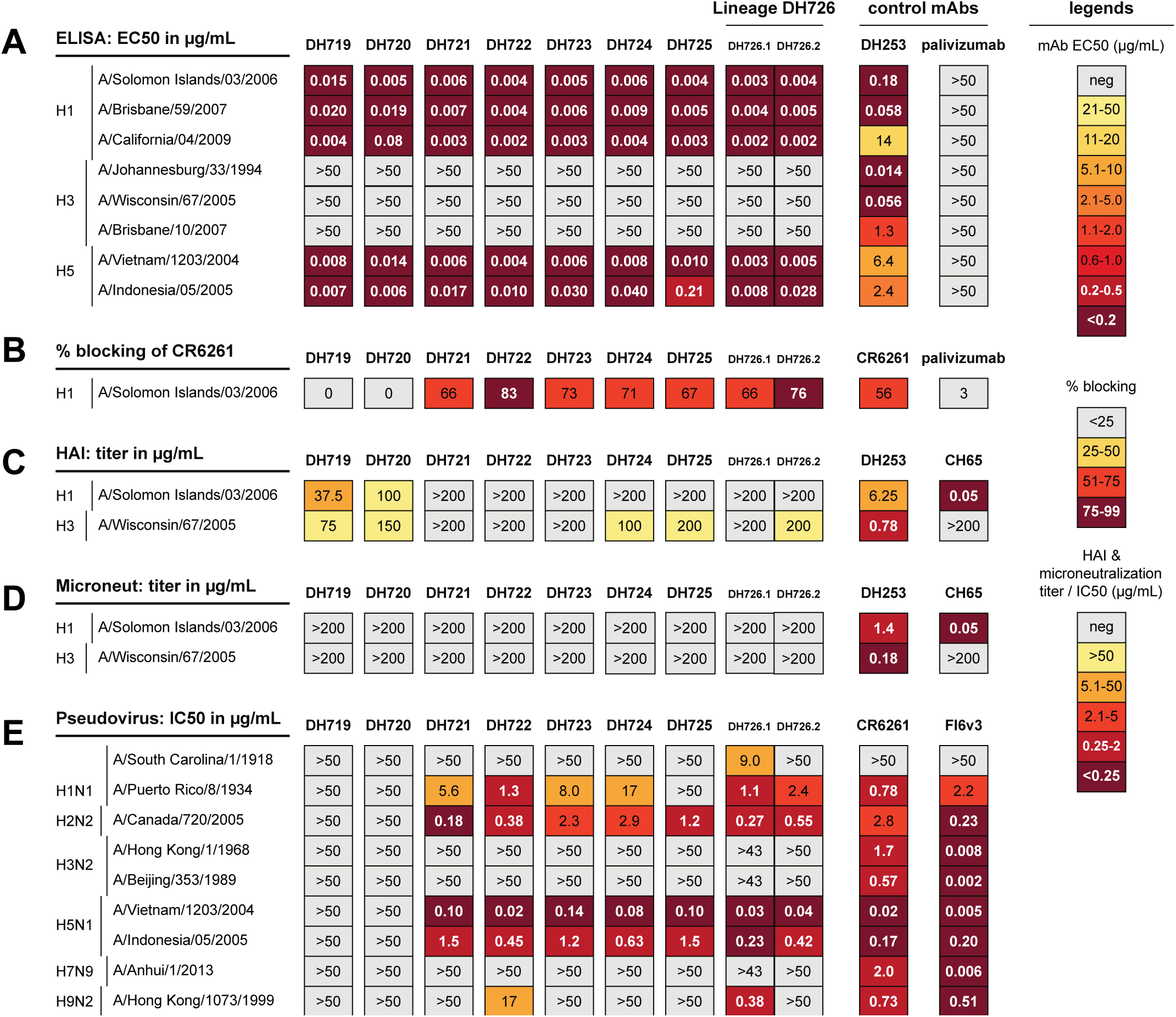
Characterization of HA-reactive mAbs. Heat map displays of mAb binding to HA in ELISA (A), blocking of CR6261 with the tested mAb at 1.1 µg/mL (B), HAI (C), microneutralization (D), and pseudovirus neutralization (E). Values are color coded relative to the legends on the right. ELISA binding shown as half maximal effective concentration (EC_50_). HAI and microneutralization shown as endpoint titer; pseudovirus neutralization shown as half maximal inhibitory concentration (IC_50_). Control mAbs are specific for each assay and are labeled.

**Table 1.** Characteristics of HA-reactive rhesus mAbs.

| Antibody name | Isotype / light chain type | Gene usage (V/(D)/J)<br>(heavy chain / light chain) | CDR3 length<br>(amino acids) | Mutation frequency |
| --- | --- | --- | --- | --- |
| DH719 | IgG | 4~F*01 / 2~C16*01 / 4~1*01 | 9 | 5.60% |
|  | lambda | 7~A*01 / 1~LC1*01 | 10 | 24.30% |
| DH720 | IgG | 4~L*03 / 3~12*01 / 4~1*01 | 16 | 2.60% |
|  | lambda | 2~D*02 / 2~2*01 | 10 | 2.80% |
| DH721 | IgG | 3~J*02 / 3C~12*01 / 5~1*01 | 20 | 5.70% |
|  | kappa | 2~S17*01 / 2~4*01 | 9 | 2.00% |
| DH722 | IgG | 3~J*02 / 3C~12*01 / 4~1*01 | 20 | 8.40% |
|  | kappa | 3~S11*01 / 2~4*01 | 8 | 4.20% |
| DH723 | IgG | 3~J*02 / 3~21*01 / 5~1*01 | 20 | 7.70% |
|  | kappa | 2~S16*01 / 1~5*01 | 9 | 3.90% |
| DH724 | IgG | 3~J*01 / 3C~12*01 / 5~1*01 | 20 | 6.40% |
|  | kappa | 1~S14*01 / 4~LC1*20 | 9 | 6.60% |
| DH725 | IgG | 3~J*01 / 2C~9*01 / 4~1*01 | 20 | 5.00% |
|  | lambda | 2~E*01 / 2~2*01 | 10 | 2.80% |
| DH726.1<br>* | IgG | 3~J*01 / 2~C13*01 / 5~1*01 | 20 | 7.70% |
|  | kappa | 1~20*01 / 4~LC1*20 | 9 | 5.80% |
| DH726.2<br>* | IgG | 3~J*01 / 4C~17*01 / 5~1*01 | 20 | 8.00% |
|  | kappa | 1~11*01 / 4~LC1*20 | 9 | 4.10% |

\*DH726.1 and DH726.2 are clonally related.

None of these rhesus mAbs neutralized A/Solomon Islands/03/2006 (H1N1) or A/Wisconsin/67/2005 (H3N2) in a standard assay with egg-grown influenza viruses (Figure 3D), but in a pseudovirus neutralization assay using HAs expressed in mammalian cells via a lentiviral vector, the seven CR6261-blocking antibodies neutralized multiple strains including an H5 (Figure 3E).

### Cryo-EM structure of DH726.1 Fab in complex with H1/SI FLsE

Single-particle cryo-EM of H1 A/Solomon Islands/03/2006 full-length solubilized ectodomain (FLsE) trimer in complex with DH726.1 Fab showed that most particles had C3 symmetry (Table S1, Figure S2, Figure S3A and S3B). A three-dimensional reconstruction of the complex (from ∼180,000 contributing particles), refined to ∼3.0Å resolution in CryoSPARC (*27, 28*) (Supplemental Figure 3C-E), revealed that DH726.1 binds the HA stem exclusively through heavy chain (HC) contacts (724 Å^2^ buried surface area) (Fig 4A). HC complementarity determining region 2 (HCDR2) forms a cleft to accommodate an HA β-strand (S361 to A365), allowing formation of a set of hydrogen bonds between the Fab and HA (Fig. 4B and Table S2). The tip of HCDR3 is a hydrophobic loop (I100-F100a-G100b-L100c-V100d-I100e) that fits into a hydrophobic pocket formed by HA G349, W350, I374 and I377 (Figure 4C). HC F100a anchors this interface.

**Figure 4.**
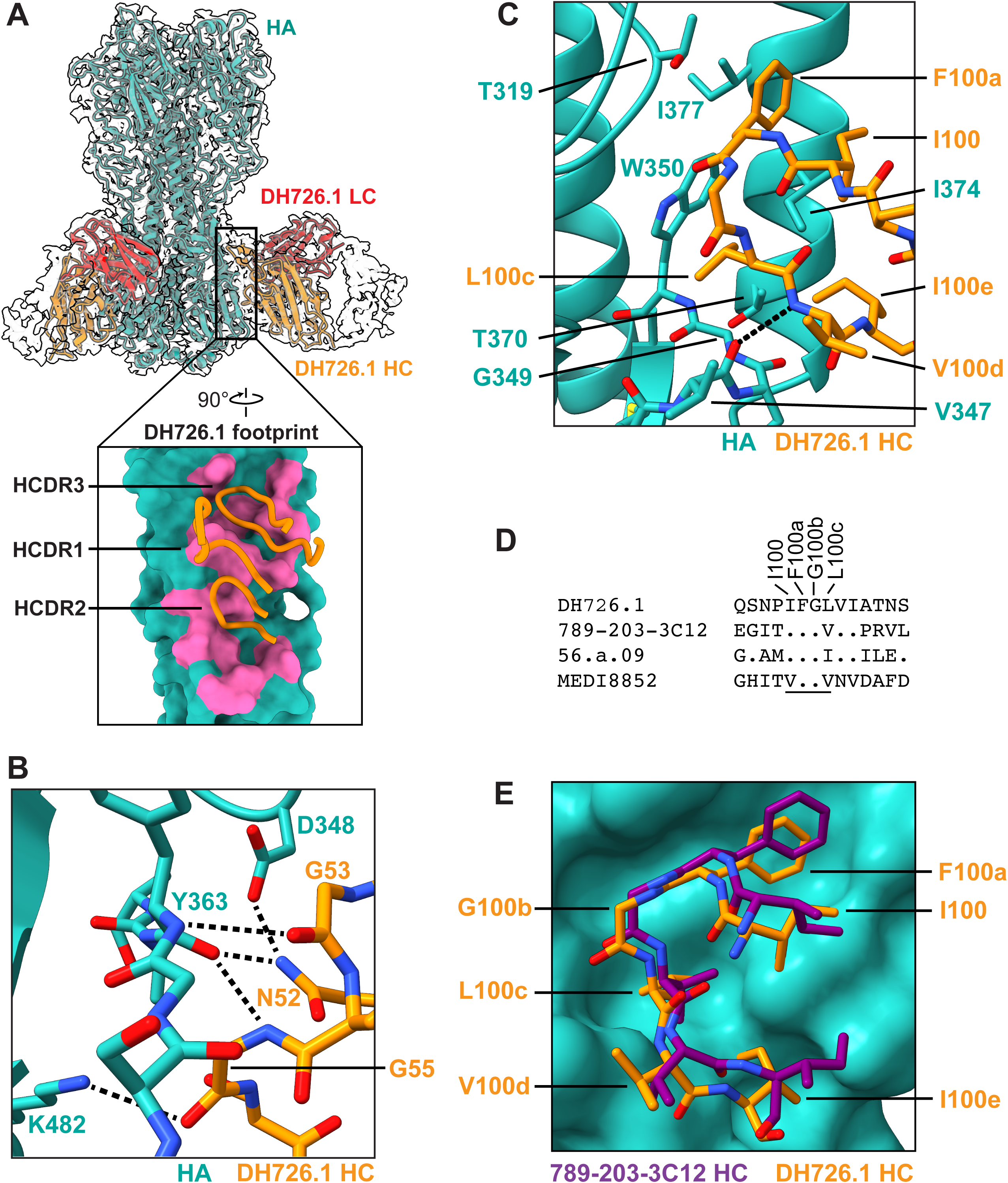
Structural analysis of DH726.1 Fab bound to HA reveals a common solution for binding a hydrophobic groove in the HA stem. (A) A 3Å resolution cryo-EM map for DH726.1 Fab bound to H1/SI06, with the resulting atomic model (colored ribbons) superposed. The inset shows the HA residues (pink surface) buried by DH726.1 HCDRs (orange loops). The turquoise surface depicts HA residues not buried by DH726.1 on a single HA protomer. (B) The interface between DH726.1 HCDR2 (orange sticks) and HA (turquoise) (Kabat convention). Hydrogen bonds are shown as black dashed lines. (C) The interface between DH726.1 HCDR3 (orange sticks) and HA (turquoise). A main-chain hydrogen-bond between HC V100d and HA V347 is shown as a black dashed line. Extensive non-polar interactions include those of HC F100a with HA T319, W350, I374, and T378 and of L100c, V100d, and I100e with various HA residues. (D) A multisequence alignment of amino acids from the HCDR3s of four antibodies. Residue numbers denote positions in DH726.1. The underlined residues make up a conserved hydrophobic motif that fits into the hydrophobic pocket of the HA stem. (E) Superposition of the atomic models of DH726.1 (orange sticks) and 789-203-3C12 (purple sticks) bound to HA (turquoise surface). The hydrophobic HCDR3 loops of the two mAbs have similar structures.

Like DH726.1, many antibodies that bind the HA stem have HCDR3s that include a hydrophobic loop with a Phe-Gly motif (Figure 4D) (*12, 29*). Examples include mAb 789-203-3C12 (from a cynomolgus macaque immunized with an experimental HA immunogen) (*12*), a multidonor class of V_H_6-1+D_H_3-3-encoded human antibodies (represented by mAb 56.a.09) (*29*), and human mAb MEDI8852 (*30*). Superposition of the structures of H1/SI06-bound DH726.1 and 789-203-3C12 showed that the HCDR3s of the two antibodies contact HA in similar, though subtly distinct, ways (Figure 4E). Thus, our HA stem-directed rhesus mAb has similarities to other stem mAbs from primates, and suggests that stem-directed antibodies can derive from diverse immune stimuli.

### Antibody-dependent cellular cytotoxicity (ADCC) activity of rhesus stem-directed mAbs

We tested whether DH726.1 and other mAbs could mediate ADCC, using a reporter assay with Flp-In T-Rex 293 cells expressing HA as target. DH722, DH726.1, and DH726.2 all induced ADCC against H1 A/New Caledonia/20/1999, H1 A/California/04/2009, and H5 A/Vietnam/1203/2004 (Figure 5A). Consistent with the lack of binding to H3 (Figure 3), the antibodies had no ADCC activity against H3 A/Aichi/2/1968 (Figure 5B). We also tested DH726.1 expressed in a murine backbone, and observed that DH726.1_Mu_IgG2a triggered FcγRIV signaling in an ADCC assay using target cells expressing H1 A/California/04/2009 or H1 A/Solomon Islands/03/2006, while DH726.1_Mu_IgG1 failed to induce FcγRIV signaling (Figure 5C). Control mAb S5V2-29_Mu_IgG2c also activated the FcγRIV signaling-cascade to H1 A/California/04/2009 targets, as reported (*31*).

**Figure 5.**
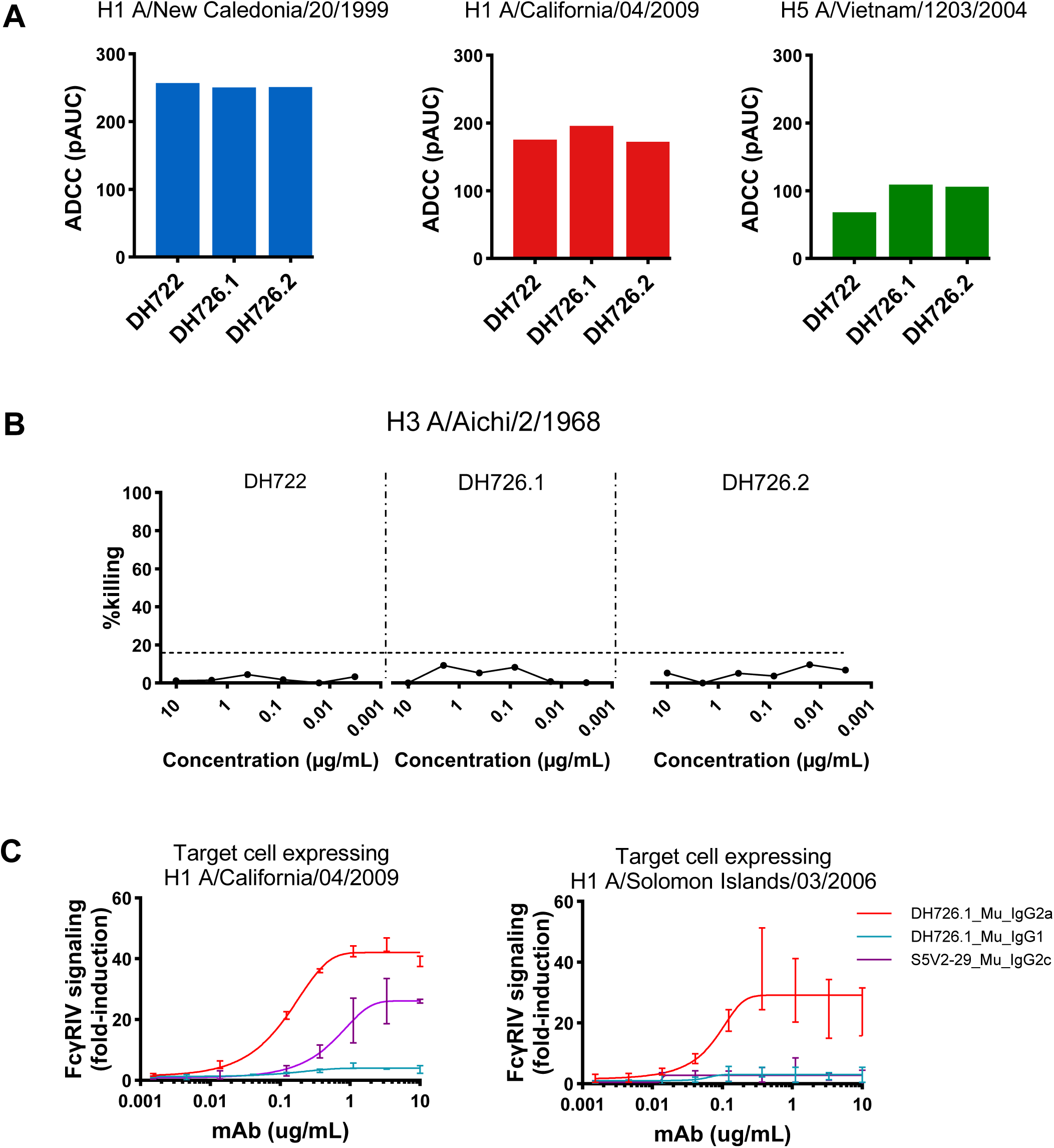
ADCC activity of rhesus stem-directed mAbs. (A) Antibody-dependent cellular cytotoxicity (ADCC) activity of DH722, DH726.1, and DH726.2 was measured using Flp-In T-Rex 293 cells expressed H1 A/New Caledonia/20/1999, H1 A/California/04/2009, and H5 A/Vietnam/1203/2004. Area under the curve (AUC) was shown. (B) No ADCC activity of mAbs was detected against H3 A/Aichi/2/1968. (C) Results of an *in vitro* ADCC proxy assay. Mouse FcγRIV-expressing effector cells and HA-expressing target cells were cocultured in the presence of serially diluted mAbs. S5V2-29_Mu_IgG2c was used as control. FcγRIV activation was measured as luminescence output.

### DH726.1 protects mice against lethal influenza challenge

Stem-directed mAbs are weakly neutralizing (12, 13) but can access the HA stem in assays using pseudotyped viruses on which HA packing is less dense (*32*). *In vivo* protection by stem-directed antibodies is Fc-receptor (FcR)-mediated (*33, 34*), so we tested whether a murinized version of DH726.1 protects mice against lethal infection. Mice were given 6 mg/kg of mAb (DH726.1_Mu_IgG2a, FI6_Mu_IgG2c (*35*), or HC19_Mu_IgG2c (*36*)) before challenge with 5× LD50 of mouse-adapted A/California/04/2009 (H1N1). All mice treated with H3 antibody HC19_IgG2c lost 25% body weight by 5 to 8 d postinfection and were euthanized, but mice treated with DH726.1_Mu_IgG2a or FI6_Mu_IgG2c were protected (Figure 6A). When mAbs DH726.1 or FI6 were given 2 days after infection as treatment, only one animal in the DH726.1_IgG2a-treated group survived (Figure S4A), indicating that the antibody needed to be present earlier to provide protection. We assayed for virus shedding by oral swabs on days 3 and 6 after infection and found shedding was similar for all groups, regardless of the mAb used or whether it was given before or after infection (Figure S4B).

**Figure 6.**
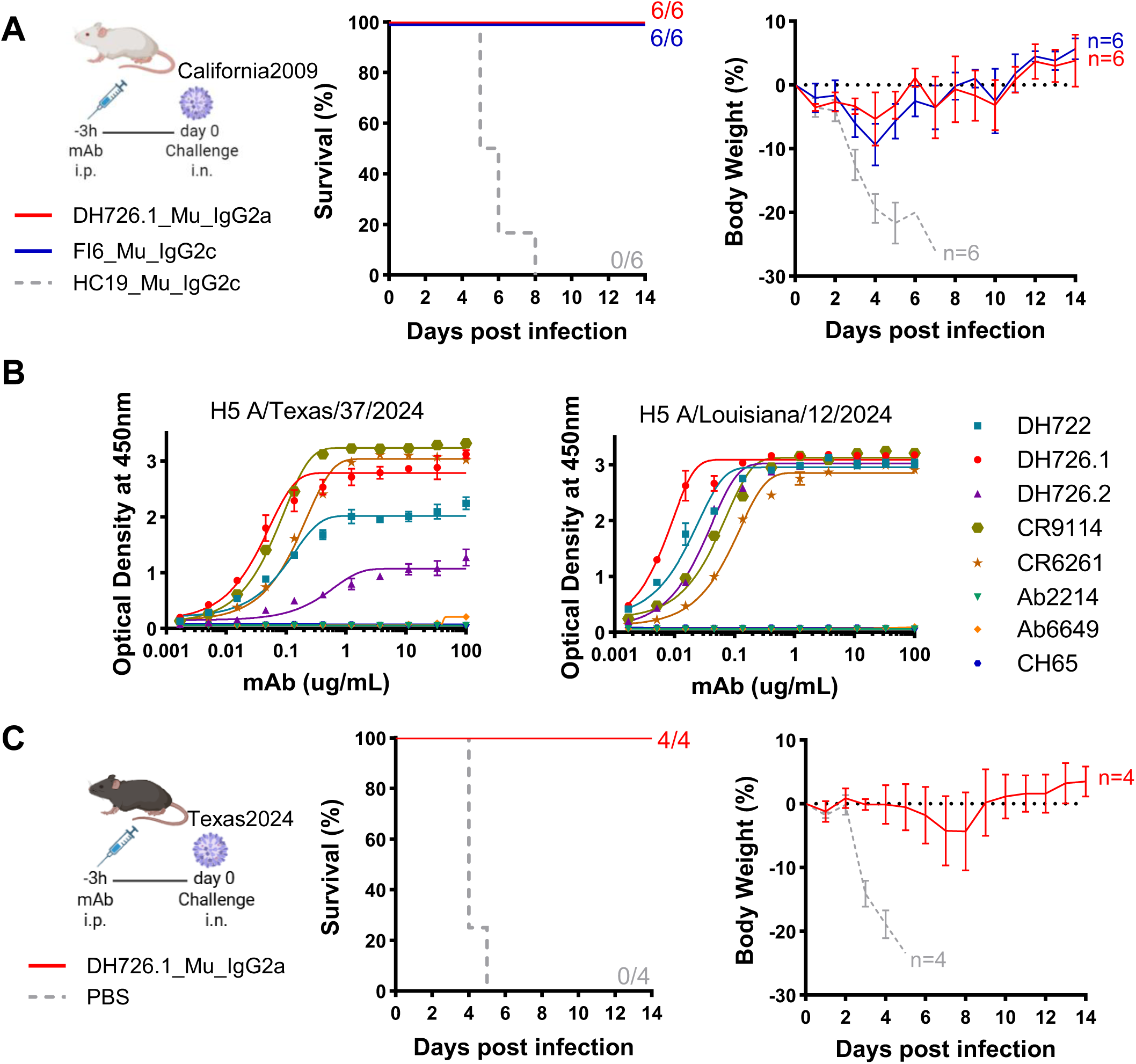
Protection by DH726.1 against lethal H1N1 and H5N1 challenge. (A) Pre-exposure prophylaxis experiment. BALB/c mice were treated with 6mg/kg of mAbs intraperitoneally 3 h prior to the intranasal virus challenge. Virus dose was 5x LD_50_. Group size was 6. Percent survival (left) and percent body weight change (right) are shown. Error bars indicate SD. (B) ELISA binding curves of CR6261-like mAbs mAbs against recombinant H5 protein of A/Texas/37/2024 and A/Louisiana/12/2024. Data are plotted as mean. (C) Pre-exposure prophylaxis experiment. C57BL/6 mice were treated with 5mg/kg of mAb or same volume of PBS (20uL) intraperitoneally 3 h before infection with 20PFU A/Texas/37/2024 virus. Group size was 4. Percent survival (left) and percent body weight change (right) are shown. Error bars indicate SD.

Since our rhesus mAbs bound to all group 1 HAs that we tested (Fig. 3A), we hypothesized they may bind more contemporary group 1 HAs. We tested DH722, DH726.1, and DH726.2 in ELISA against recent HAs from H5 A/Texas/37/2024 and H5 A/Louisiana/12/2024 (Figure 6B). Our rhesus mAbs elicited by ordinary HA immunization bound these HAs, as did stem mAbs CR9114 and CR6261; mAbs directed to other HA epitopes such as the lateral patch (Ab6649), apex/RBS (CH65), and trimer interface (Ab2214) did not (Figure 6B). We then tested whether DH726.1_IgG2a would protect treated mice in a lethal A/Texas/37/2024 (H5N1) infection model, finding that all treated mice survived with minimal weight loss and clinical signs (Figure 6C and Figure S4C). We conclude that mAbs elicited by ordinary vaccination can have the ability to protect against a heterologous influenza strain.

## Discussion

Influenza vaccination remains one of the best ways to prevent influenza spread and protect against serious complications (*37*). Antigenic drift and shift can result in vaccine mismatch (*38, 39*), so many vaccine design efforts have focused on targeting relatively conserved influenza epitopes (*40–42*), including the HA stem. It is unclear whether special vaccine designs are necessary to elicit such responses. In this study, we identified and characterized HA stem-directed mAbs elicited in rhesus macaques using an unremarkable HA immunogen. The stem-directed mAbs we isolated have potency and binding specificity similar to those from humans and share features of human mAbs isolated in other studies (*43*). Unlike mAbs against other pathogens like HIV-1 (*44*), these stem-directed antibodies do not appear to have features that make them difficult to elicit by vaccination.

Animal models are useful but do not always recapitulate what is seen in humans. Rhesus macaques are advantageous for vaccine studies since they are similar to humans at the genetic level (*45*). Human HA stem-directed antibodies often use V_H_1-69 gene segments (*22, 43*) a predominance due to the presence of germline-encoded Phe residues in HCDR1 and HCDR2 (*11*), which give antibodies using V_H_1-69 a “head start” in the affinity maturation process. Rhesus macaques lack an ortholog of the human V_H_1-69 gene segment, but the two species do share similar V_H_3 gene segments, and humans have also been shown to make stem-directed antibodies using V_H_3 (*43*). Other vaccine studies in NHPs using headless HA stabilized-stem ferritin nanoparticle or HA stem-ferritin immunogens elicited stem antibodies (*12, 46*), showing the utility of the model.

The stem-directed V_H_3-encoded rhesus antibodies we isolated were elicited by an ordinary H1 HA protein, but they exhibited intra-group cross-reactivity to H5 proteins. These rhesus mAbs did not neutralize in a traditional microneutralization assay with egg-grown influenza viruses, but they could neutralize multiple strains, including H5N1, in a pseudovirus-based assay. These results are consistent with the inaccessibility of the HA stem on mature influenza virions (*47–49*), because of tight packing of the HA trimer on the virion surface. The lentivirus-based pseudovirions probably have much less closely spaced surface HAs. Lack of neutralizing activity against authentic influenza virus implies that this group of antibodies are unlikely to prevent influenza infection. Because these antibodies mediated ADCC, and protected mice against lethal challenge with A/California/04/2009 (H1N1) and A/Texas/37/2024 (H5N1), they may have public-health benefits. The H5N1 strain we used was derived from a human infected during the recent dairy farm outbreaks (*50*), suggesting that antibodies of this type may have helped prevent severe disease among humans exposed to these zoonotic strains. Thus, a class of antibodies easily elicited by an unremarkable immunogen suggests that routine vaccination may be more protective than previously appreciated, even if it cannot prevent infection.

## Resource availability

## Lead contact

Further information and requests for resources and reagents should be directed to and will be fulfilled by Dr. M. Anthony Moody.

## Materials availability

All unique/stable reagents generated in this study are available from the lead contact with a completed materials transfer agreement.

## Data and code availability

1. All data reported in this article will be shared by the lead contact upon request.
2. The map of the cryo-EM reconstruction of DH726.1 Fab bound to H1/SI are available at the Structural Bioinformatics Protein Data Bank (accession number 9E6J) and the Electron Microscopy Data Bank (accession number EMD-47570).

### Acknowledgments

We thank Charles E. McGee, Nathan A. Vandergrift, Thaddeus C. Gurley, John F. Whitesides, Ashley A. Allen, Lauren E. Stiegel, and Christopher J. Sample for expert technical support. Support for this work was provided by grants from the NIH NIAID, including the Division of Microbiology and Infectious Diseases P01-AI089618 (to SCH and MAM) and the Division of AIDS, Center for HIV/AIDS Vaccine Immunology (CHAVI) U19-AI067854 (to BFH). The graphical abstract was created in BioRender.

## Author Contributions

S.G., B.F.H., and M.A.M. designed research; S.G., J.F., K.L., S.M.V., D.M., J.P., T.A.V., D.J.M, R.P., L.L.S., R.M.S., S.H., J.D., M.A.S. performed research; K.W. contributed new reagents/analytic tools; S.G., J.F., S.S., C.D.L., N.S.H., H.M.Y., B.S.G., T.B.K., H.L., A.G.S., G.F., B.F.H., S.C.H., M.A.M. analyzed data; and S.G., J.F., and M.A.M. wrote the paper.

---

**Supplemental Figure 1.** (A) Trimeric HA constructs were formulated as B cell tetramers and tested on antibody coated beads. (B) Frequency of VH and VL gene somatic mutation in isolated memory B populations.

**Supplemental Figure 2. Schematic depicting the strategy for processing cryo-EM data.**

**Supplemental Figure 3.** (A) Representative micrograph of DH726.1 Fab bound to H1 A/Solomon Islands/03/2006, embedded in vitreous ice (scale bar = 200 Å), low pass filtered for clarity. (B) Selected 2D class averages of DH726.1 Fab:HA complexes. (C) 3-D reconstruction of three DH726.1 Fabs bound to HA trimer, filtered and colored by local resolution. (D) Gold-standard Fourier shell correlation (GSFSC) curves from CryoSPARC. (E) Viewing direction distribution plot.

**Supplemental Figure 4. Mouse challenge study.** (A) Post-exposure treatment experiment. BALB/c mice were infected with virus prior to mAb administration (5 mg/kg) at 2 days post-infection. Virus dose was 5x LD50. Group size was 6. Percent survival (left) and percent body weight change (right) are shown. (B) Viral titers from oral swabs collected on days 3 and 6 in mice treated with either therapeutic or prophylactic groups. (C) The clinical disease scores for the A/Texas/37/2024 virus challenge study. Clinical score metrics: Score 0: Normal, no signs of illness; Score 1: Mild illness — the mouse appears slightly unwell (e.g., mildly hunched posture) but remains active and moves around the cage; Score 2: Moderate illness — clearly sick (hunched posture, ears back), but still mobile with or without stimulation; Score 3: Severe illness — very sick, hunched, sluggish, and largely immobile, even when provoked.

## STAR Methods

### Immunization of rhesus macaques

The vaccine regimen for this study is shown in Figure 1A. In brief, six male, adult rhesus macaques were immunized with HA from H1 A/Solomon Islands/03/2006 at a dose of 100 μg per injection adjuvanted in a 1:1 ratio with MF59 at weeks 0, 8, 12, 16, and 20 (1). Peripheral blood was obtained at each immunization and 2 weeks after each immunization. Peripheral blood was processed for serum or plasma and peripheral blood mononuclear cells (PBMCs); all samples were aliquoted and maintained at −80°C (serum / plasma) or in liquid nitrogen vapor phase (PBMCs) for further analysis.

All rhesus macaques were housed at New England Primate Research Center, Southborough, MA and treated in accordance with the regulation of Association for Assessment and Accreditation of Laboratory Animals with the approval of the Animal Care and Use Committees of the National Institutes of Health and Harvard Medical School, Cambridge, MA.

## METHOD DETAILS

### ELISA assays of rhesus plasma and mAbs

Plasma from each RM was heat inactivated at 56°C / 5 min before assay by ELISA. Influenza HA antigens were from the following strains: H1 A/Solomon Islands/03/2006, H1 A/Brisbane/59/2007, H1 A/California/04/2009, H3 A/Wisconsin/67/2005, H5 A/Vietnam/1203/2004, and H5 A/Indonesia/05/2005. Antigens were coated at 4°C on 384-well Costar high binding ELISA plates at 30 ng per well. After overnight incubation, ELISA plates were blocked by superblock (40 g whey protein, 150 mL goat serum, 5.0 mL Tween20, 0.5 g NaN_3_, 40 mL of 25× PBS brought to 1 L with deionized water) for 1 h at room temperature. Plasma samples were diluted 3 fold for 12 dilutions in Superblock buffer and added at 10 µL per well. After 1 h room temperature incubation, plates were washed twice with PBS containing 0.1% Tween20. Horseradish peroxidase-conjugated goat anti-monkey secondary antibodies (Rockland) were 1:10000 diluted in superblock and added at 10 µL per well for a 1 h incubation. Plates were washed 4 times before 20 µL of SureBlue Reserve TMB-1 Solution (KPL) was added. After 15 min incubation, the reaction was stopped with 20 µL of 1% HCl. Plates were read by a Spectramax 384plus plate reader (Molecular Devices) at 450 nm. Area under the curve (AUC) was calculated using the trapezoidal method. Control monoclonal antibodies (mAbs) included stem-directed influenza mAb CR6261 (2, 3), receptor binding site (RBS) mAb CH65 (4), cross-reactive mAb DH253 (5), stem-directed mAb FI6v3 (6), and anti-respiratory syncytial virus mAb palivizumab (7).

Competitive inhibition ELISAs were performed as described (8) using a biotinylated or HRP-conjugated version of influenza stem-directed mAb CR6261 (2, 3) as the mAb being competed for binding to HA H1 A/Solomon Islands/03/2006, H5 A/Vietnam/1203/2004, or H1 A/California/07/2009. Plasma samples were tested at 1:50 dilution; mAbs were tested over 12 five-fold dilutions starting at 100 µg/mL.

For V_H_ / V_L_ transient transfection supernatants, screening was performed as described (9, 10) against HA H1 A/Solomon Islands/03/2006. The mAb concentration in each supernatant was quantified using a capture ELISA method as described (11); known concentrations of mAb 2F5 were used to generate the standard curve. Supernatants were undiluted for ELISA binding screening. For mAbs produced at larger scale and purified, a panel of HAs were tested including H1 A/Solomon Islands/03/2006, H1 A/Brisbane/59/2007, H1 A/California/04/2009, H3 A/Johannesburg/33/1994, H3 A/Wisconsin/67/2005, H3 A/Brisbane/10/2007, H5 A/Vietnam/1203/2004, H5 A/Indonesia/05/2005, H5 A/Texas/37/2024, and H5 A/Louisiana/12/2024. Concentrations of mAbs were determined by Nanodrop and tested using 3-fold dilutions 12 times starting at 100 µg/mL; ELISA was performed as described above.

### Influenza microneutralization and hemagglutination inhibition assay of plasma and mAbs

The microneutralization assay was based on a standard published protocol (12). Influenza stocks were grown in embryonated eggs and were titrated for hemagglutination units on turkey red blood cells. Madin-Darby canine kidney (MDCK) epithelial cells were cultured as monolayers in 96-well culture plates at 37°C in 5% CO_2_, followed by the addition of mixtures of serial dilutions of plasma or purified mAbs and 100 TCID_50_ of the appropriate infectious influenza virus stock. Plates were incubated at 37°C in 5% CO_2_ for 18 h prior to assay. Final assay of the infectious cultures was performed by using fluorescence (for virus constructs containing fluorescent reporter genes); by ELISA using mouse mAb anti-influenza A nucleoprotein (BEI resources) at 1:4000 dilution, followed by HRP-conjugated Goat anti-mouse IgG (KPL) at 1:2,500 dilution (according to CDC SOP); or by testing of the resulting virus cultures by hemagglutination inhibition (HAI).

HAI was based on a standard published protocol (12). To perform HAI assays, serial dilutions in PBS of plasma or transfected cell supernatants were placed into 96-well plates and were mixed with an equal volume of washed turkey red blood cells (0.5%) and incubated at room temperature for 30 min before hemagglutination was read directly from the wells.

### Single-cell sorting by flow cytometry

Preformed conjugates for antigen-specific B cell sorting were made as described (13) using biotinylated HA protein coupled to streptavidin conjugates AlexaFluor 647 (Life Technologies) or BV421 (Biolegend). Antigen-specific probes were made using trimeric HAs from H1 A/Solomon Islands/03/2006, H1 A/California/04/2009, H3 A/Wisconsin/67/2005, and H5 A/Vietnam/1203/2004. Probes were tested at multiple concentrations for binding to a panel of mAbs attached to polystyrene beads (Spherotech); flow data were acquired on a BD LSRII flow cytometer (BD Biosciences) and analyzed using FlowJo (Treestar).

Isolated cells were stained with a panel of fluorochrome antibody conjugates and reagents to identify antigen-specific memory B cells from the PBMCs. The panel consisted of Aqua Vital dye (Life Technologies); CD3 (peridinin chlorophyll protein-cyanine dye 5.5 [PerCP-Cy5.5]), CD16 (phycoerythrin [PE]-Cy7), CD20 (FITC) (BD Biosciences); CD14 (Brilliant Violet [BV]570), CD27 (allophycocyanin [APC]-Cy7) (Biolegend); and IgD (PE) (Southern Biotech). Cell isolation was performed using a BD FACS Aria2 (BD Biosciences) cell sorter. Memory B cells were defined as viable CD3^−^CD14^−^CD16^−^CD20^+^IgD^−^; antigen-specific memory B cells positive for probes in both colors were sorted as single cells into 96-well plates containing 20 μL/well reverse transcriptase (RT) buffer (Invitrogen) as described (9). Sorted plates were frozen immediately and maintained at −80°C before RT/PCR. Flow cytometry data was analyzed using FlowJo (Treestar).

### PCR amplification of immunoglobulin V_H_ and V_L_ genes

The variable heavy chain (V_H_) and light chain (V_L_) genes of sorted single memory B cells were amplified as described (9, 14). A 65°C 5 min pre-RT incubation was performed by adding 0.1 μM of mixture of Ig constant region primers (including IgA, IgM, IgD, IgG and IgE) to each well. Samples were chilled on ice and RT buffer was added following a 50°C / 45 min, 55°C / 15 min RT/PCR reaction. First round PCR (PCRa) of heavy, kappa and lambda chains was then performed using the cDNA product of RT/PCR. Each 50 µL reaction contained 5 µL of cDNA, 0.6 µL of HotStar taq (Qiagen), 5 µL of PCR Buffer (Qiagen), 10 µL of Q Buffer (Qiagen), 0.4 µL of 25 mM dNTPs (Qiagen), 25 mM MgCl_2_ (1 µL for IgH, 2 µL for Igκ and 3 µL for Igλ) and 0.125 µM of IgH, Igκ, or Igλ variable region primer mixtures. PCRa reaction conditions were as follows: 95°C / 5 min, [94°C / 30 sec, 62°C (for IgH) or 64°C (for Igκ / λ) / 45 sec, 72°C / 90 sec] for 35 cycles, 72°C / 7 min, then hold at 10°C.

PCRa products were used as templates for nested PCR (PCRb). Each 50 µL reaction contained 3 µL of PCRa product, 0.6 µL of HotStar taq (Qiagen), 5 µL of PCR Buffer, 10 µL of Q Buffer, 0.4 µL of 25 mM dNTPs, 25 mM MgCl_2_ (3 µL for IgH, 2 µL for Igκ/λ) and 0.125 µM of IgH, IgK, or Igλ internal primers. PCRb reaction conditions were the same as PCRa above. PCRb products were then analyzed by 1.2% SYBER® Safe E-Gels® (Invitrogen). Wells with bands indicating a PCR product were purified and sequenced. Sequencing results were analyzed by SoDA (15) and ARPP (16) software to infer VDJ arrangements.

Analysis of rhesus genes and clonal lineages was performed as described (14, 16, 17). Clonal lineage membership was determined using Cloanalyst (16, 17), which takes into account V and J gene usage, CDR3 length, and overall sequence similarity. Cloanalyst reconstructs lineage relationships based on maximum posterior probability trees and groups sequences accordingly.

### Recombinant mAb expression

Initial screening of isolated mAbs was performed using transient transfection as described (9, 14). To produce the transfection construct, 2 µL of V_H_/V_L_ PCRb product, 1 µL of KOD polymerase (Novagen), 5 µL of KOD Buffer, 5 µL of dNTPs, 3 µL of 25 mM MgSO_4_, 1 µL of CMV-262, 1 µL of 1822BGH (IgH) or BGH-1235 (IgK/λ), 2 µL of rhesus CMV_P DNA fragments, 2 µL of reversed DNA primers for either IgH or IgK/λ were brought to 50 µL with H_2_O. PCR conditions were as follows: 95°C / 2min, [95°C / 20 sec, 62°C / 12 sec, 70°C / 60 sec] for 30 cycles, then hold at 4°C. PCR products were purified by MinElute 96UF plates and analyzed by QIAxcel (Qiagen) before transient transfection.

Transient transfection was performed using 293T cells cultured in 6-well plates. About 1 µg of heavy and light chain recombinant linear fragments were co-transfected by Effectene (Qiagen) when cells cultured in 10% FBS supplemented DMEM were 80-90% confluent. Cell medium was changed to 2% FBS / DMEM before adding the transfection mixture. After three days of incubation at 37°C / 5% CO_2_, supernatants were harvested for analysis.

For large-scale production of mAbs, isolated V_H_ and V_L_ genes were cloned into pcDNA3.1^+^ plasmid containing the CMV promoter and IgG constant regions (GeneScript). Plasmids were amplified by plasmid Plus Mega kit (Qiagen) and concentrated. V_H_ and V_L_ plasmids were co-transfected into Expi293F cells by PEI transfection reagent (EMD Millipore). Medium was changed to FreeStyle^TM^ 293 Expression Medium (ThermoFisher) at 6 h after transfection. Cells were cultured on a 120 rpm shaker at 37°C / 8% CO_2_. After 5 days of incubation, supernatants were harvested and mAbs purified by Protein A agarose beads (Thermo). SDS-PAGE and Western blots under reducing and nonreducing conditions were used to analyze antibody quality (9).

### Influenza pseudovirus neutralization assay

The pseudotyped influenza neutralization assay was performed in 293A cells using luciferase encoding lentiviruses pseudotyped for influenza HA as described previously (18). HA sequences used to generate pseudoviruses were derived from: Group 1 [A/South Carolina/1/1918 (H1N1), A/Puerto Rico/8/1934 (H1N1), A/Canada/720/2005 (H2N2), A/Vietnam/1203/2004 (H5N1), A/Indonesia/05/2005 (H5N1), and A/Hong Kong/1073/1999 (H9N2)] and Group 2 viruses [A/Hong Kong/1/1968 (H3N2), A/Beijing/353/1989 (H3N2), and A/Anhui/1/2013 (H7N9)]. Briefly, monoclonal antibodies were serially diluted and incubated with pre-titrated HA pseudotyped viruses before addition to 293A cells. Pseudovirus neutralization shown as half maximal inhibitory concentration (IC_50_).

### Antibody-dependent cellular cytotoxicity reporter assay

To generate target cell lines, single copies of plasmids containing Influenza HA genes connected to a Renilla luciferase gene by an IRES2 sequence (HA-IRES2-LucR) and an FRT site were integrated at the FRT site of Flp-In T-Rex 293 cells by dual transfecting with a plasmid containing the Flp Recombinase gene as per manufacturer’s recommendations (Thermofisher). The day prior to the antibody-dependent cellular cytotoxicity (ADCC) assay, target cells were activated to express HA and Renilla Luciferase by suspending cells in DMEM containing 10% Tet-system approved fetal bovine serum (Gibco), 15µg/mL Blasticidin (Santa Cruz), 100µg/mL Hygromycin (Gibco) and 2µg/mL Doxycycline (Santa Cruz) and plated into opaque tissue culture treated white 96-well plates (Corning) at 2.5×10^4^ cells/well. NK92 cells expressing human CD16 receptors containing the high affinity V158 SNP (hCD16.NK92; Conkwest Inc.) were concentrated to 1×10^6^ cells/mL in Myelocult media (Stemcell) containing 100IU/mL IL-2. On the day of the assay, hCD16.NK92 cells were counted, assessed for viability, and the concentration adjusted to 1×10^6^ cells/mL (5×10^4^ cells/well; effector/target ratio ∼ 1:1) in AIM V media (Gibco) containing 100IU/mL IL-2. mAbs were serially diluted five-fold in AIM V media with a final concentration of 10µg/mL. hCD16.NK92 cells and antibody dilutions were added to plates, centrifuged at 300 g for 1 min after 30 min, and then incubated for 5.5 h at 37°C, 5.5% CO_2_ to allow antibody-mediated cell lysis to proceed. After 5.5 h, ViviRen substrate (Promega) was diluted 1:250 in AIM V media and added 1:1 to the assay wells. The substrate generates luminescence only in live cells; not in dead or lysed cells. The final readout was the luminescence intensity generated by the presence of residual intact target cells that have not been lysed by the effector population in the presence of ADCC-mediating antibodies. The percentage of specific killing was calculated after subtracting background luminescence intensity of unactivated target cells from all wells using the formula

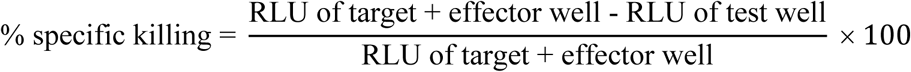

In the analysis, the relative light units (RLU) of the target + effector wells represent lysis by hCD16.NK92 cells in the absence of any source of antibody. The anti-RSV antibody, Synagis, was used as a negative control and the anti-Influenza HA stem antibody, CR9114, was used as a positive control.

### Recombinant HA expression and purification

Expi293F cells (Homo sapiens; Thermo Fisher) were cultured in Expi293 Expression Medium plus penicillin (100 units/ml) and streptomycin (100 μg/ml), at 8% CO2 with shaking. High Five cells (BTI-TN-5B1-4; Trichoplusia ni; Thermo Fisher) were maintained in ESF 921 medium (Expression Systems) at 28°C in spinner flasks in air. Cell lines were not subject to authentication.

Recombinant H1 A/Solomon Islands/03/2006 (H1/SI06) full-length soluble ectodomain (FLSE) trimer was expressed by infection of insect cells with recombinant baculovirus as described (4, 19–21). In brief, synthetic DNA encoding the HA ectodomain was subcloned into a pFastBac vector modified to encode a C-terminal thrombin or HRV3C cleavage site, a T4 fibritin (foldon) trimerization tag, and a hexa-His tag. The resulting baculoviruses produces HA trimer. Supernatant from recombinant baculovirus-infected High Five cells was harvested 72 h post-infection and clarified by centrifugation. HA was purified by adsorption to cobalt-nitrilotriacetic acid (Co-NTA) agarose resin (Takara), followed by a wash in buffer A (10 mM Tris, 150 mM NaCl, pH 7.5) plus 5 mM imidazole, elution in buffer A plus 350 mM imidazole (pH 8), and gel filtration chromatography on a Superdex 200 column (GE Healthcare) in buffer A. HA used for cryo-electron microscopy (cryo-EM) was incubated with HRV3C protease (Thermo Fisher) overnight at 4°C to remove the T4 fibritin domain and the hexa-His tag. HRV3C, tag fragments, and HA retaining the hexa-His tag were removed by a second round of adsorption to co-NTA agarose resin.

### Recombinant Fab expression and purification

DNA encoding the H- and L-chain variable domains of DH726.1 was cloned into expression vectors harboring the constant regions of human IgG1 Fab and Igκ. The Fab HC vector included a C-terminal HRV3C cleavage site and a hexa-His tag. Fab was produced by transient transfection of Expi293F cells with the Expifectamine 293 transfection kit (Thermo Fisher), according to the manufacturer’s instructions. Five days post-transfection, supernatant was harvested and clarified by centrifugation. Fab was purified by adsorption to Co-NTA resin, as described for HAs. After cleavage and removal of the hexa-His tag, Fab was dialyzed into PBS, and its concentration was determined with a NanoDrop spectrophotometer (Thermo Fisher).

### Cryo-EM sample preparation

HA A/Solomon Islands/03/2006 was incubated with 3-fold molar excess (per HA0 polypeptide chain) of DH726.1 Fab on ice for 60 min. Fab:HA complex was isolated by size-exclusion chromatography over a Superdex 200 Increase 10/300 GL column (GE Healthcare) equilibrated in PBS. Pooled fractions containing the Fab:HA complex were concentrated to 1.5 mg/ml in PBS containing 0.1% w/v octyl-beta-glucoside (ThermoFisher). 3.0 µL of sample was deposited onto Quantifoil R 1.2/1.3 400 Mesh Cu grids that had been glow discharged in a PELCO easiGLOW (Ted Pella) at 0.39 mBar, 15 mA for 30 s. Samples were vitrified in 100% liquid ethane using a Vitrobot Mark IV (Thermo Fisher Scientific), with a wait time of 2 s, blot time of 4 s and a blot force of 5 at 100% humidity.

### Cryo-EM data collection and processing

Cryo-EM data were recorded on a 200 kV Talos Arctica Microscope (Thermo Fisher Scientific) equipped with a K3 direct electron detector (Gatan) and at the Harvard Cryo-Electron Microscopy Center for Structural Biology at Harvard Medical School. Data were acquired in counting mode with the automated data collection software SerialEM (22); nine holes were visited per stage position, acquiring one movie per hole. Details of the data collection and dataset parameters are summarized in Table S1. All data processing was performed using CryoSPARC (23, 24). Details of the data processing strategy are shown in Fig. S1. Briefly, dose-fractionated images were gain-normalized, aligned, dose-weighted and summed, and then patch contrast transfer function and defocus value estimation were performed. After automated particle picking, 2D classification, and culling of junk classes, 194,142 particles were selected for ab-initio reconstruction and heterogeneous refinement of five 3D classes. The largest and best-looking class, comprising 180,474 particles, was subjected to non-uniform refinement (24) with reference-based motion correction to produce the final ∼3Å reconstruction.

### Atomic model building and refinement

UCSF ChimeraX v1.8 (25, 26) was used to fit the atomic models of one HA1+HA2 protomer of HA A/Solomon Islands/03/2006 (PDB: 5UGY) (4) and one Fab module (PDB: 6XSK) (27) to the cryo-EM map. The Fab constant domain and the N-linked glycans on HA were ignored, due to low density at these regions of the map. Coot v0.9.8.91 (28) was used to mutate the Fab to match the amino acid sequence of DH726.1. The atomic model was refined with Coot and PHENIX (29, 30). Kabat numbering was applied to the Fab residues via the Abnum server (31). ChimeraX was used for visualization and analysis of the final model. Structural biology applications used in this project were compiled and configured by SBGrid (32).

### Mouse ADCC reporter assay

The potential for mAbs to mediate ADCC activity was determined using the Mouse FcγRIV ADCC Bioassay (Promega) according to manufacturer’s instructions. Briefly, cloned HA (H1 A/California/04/2009 and H1 A/Solomon Islands/03/2006)-expressing K530 cells (target cells) were dispensed at 2.5×10^4^ cells/well into white, flat-bottom 96-well assay plates. Serially diluted mouse IgG2c Abs were added to the target cells and incubated at RT for 15 min. Effector cells expressing mouse FcγRIV were added to the wells for an effector:target ratio of 3:1, then the plate was incubated at 37°C, 5% CO_2_ for 6 h. Finally, Bio-Glo Reagent was added to the plate and incubated for 15 min at RT. Luminescence was detected with a Synergy HTX multimode plate reader (BioTek). Background signal from wells containing no cells was subtracted from the data, and fold induction was calculated as the quotient of signal in wells containing Ab divided by the signal in wells containing no Ab. Antibodies were assayed in duplicate.

### *In vivo* challenge

Female BALB/c mice at 8 weeks old (n=6 mice/mAb) were anesthetized with 5% isoflurane and intranasally infected with 5x median lethal dose (LD50) of mouse adapted A/California/04/2009 virus. Mice were given the indicated antibody at a dose of 6 mg/kg intraperitoneally at 3 h before infection (prophylaxis) or 2 days after infection (therapeutics). Weight loss was monitored daily for 14 days. The humane endpoint was defined as a weight loss of 25% from initial weight on day 0. To determine differences in viral load, oral swabs were collected at day 3, day 6 and day 9 post-infection. Subsequently, viral load was determined via plaque assay. As to A/Texas/37/2024 virus challenge, C57BL/6 mice were given the indicated antibody at a dose of 5 mg/kg (20uL) intraperitoneally at 3 h before infection with 20 PFU virus. All the animal experiments were performed in accordance with protocols proved by the Duke University Institutional Animal Care and Use Committee.

### Statistical analysis

Four-parameter logistic regression and determination of the half maximal binding concentration (EC_50_) of mAbs in ELISA was performed using the drc package in R. Area under the curve (AUC) calculations were performed using the trapezoidal method. IC_50_ values for neutralization curves were generated by GraphPad Prism. The Mann-Whitney U test and Wilcoxon matched-pairs signed-rank test were used when two groups were compared. *In vivo* challenge. Statistical significance of differences in mouse survival after lethal influenza challenge was calculated by the log-rank (Mantel-Cox) test. Statistical analysis was performed using the GraphPad Prism (version 10) software or R. All tests were two-sided, and a *P* value < 0.05 was considered statistically significant.

## References

1. World Health Organization Influenza Fact Sheet. Accessed on 3 October 2023. Available online: https://www.who.int/en/news-room/fact-sheets/detail/influenza-(seasonal).

2. J. L. Goodman, R. A. Bright, N. Lurie, H5 Influenza Vaccines—Moving Forward Against Pandemic Threats. JAMA, (2024).

3. C. M. Trombetta, O. Kistner, E. Montomoli, S. Viviani, S. Marchi, Influenza Viruses and Vaccines: The Role of Vaccine Effectiveness Studies for Evaluation of the Benefits of Influenza Vaccines. Vaccines (Basel) 10, (2022).

4. C. G. Grijalva, H. Q. Nguyen, Y. Zhu et al., Estimated Effectiveness of Influenza Vaccines in Preventing Secondary Infections in Households. JAMA Netw Open 7, e2446814 (2024).

5. C. G. Grijalva, L. R. Feldstein, H. K. Talbot et al., Influenza Vaccine Effectiveness for Prevention of Severe Influenza-Associated Illness Among Adults in the United States, 2019-2020: A Test-Negative Study. Clin Infect Dis 73, 1459–1468 (2021).

6. CDC. 2024. https://www.cdc.gov/flu/vaccines-work/effectiveness-studies.htm. Accessed August 14, 2024.

7. D. C. Ekiert, G. Bhabha, M. A. Elsliger et al., Antibody recognition of a highly conserved influenza virus epitope. Science 324, 246–251 (2009).

8. J. R. Whittle, R. Zhang, S. Khurana et al., Broadly neutralizing human antibody that recognizes the receptor-binding pocket of influenza virus hemagglutinin. Proc Natl Acad Sci U S A 108, 14216–14221 (2011).

9. S. Black, U. Nicolay, T. Vesikari et al., Hemagglutination inhibition antibody titers as a correlate of protection for inactivated influenza vaccines in children. Pediatr Infect Dis J 30, 1081–1085 (2011).

10. L. Coudeville, F. Bailleux, B. Riche et al., Relationship between haemagglutination-inhibiting antibody titres and clinical protection against influenza: development and application of a bayesian random-effects model. BMC Med Res Methodol 10, 18 (2010).

11. D. Lingwood, P. M. McTamney, H. M. Yassine et al., Structural and genetic basis for development of broadly neutralizing influenza antibodies. Nature 489, 566–570 (2012).

12. S. M. Moin, J. C. Boyington, S. Boyoglu-Barnum et al., Co-immunization with hemagglutinin stem immunogens elicits cross-group neutralizing antibodies and broad protection against influenza A viruses. Immunity 55, 2405–2418.e2407 (2022).

13. C. Dreyfus, N. S. Laursen, T. Kwaks et al., Highly conserved protective epitopes on influenza B viruses. Science 337, 1343–1348 (2012).

14. N. M. Bouvier, A. C. Lowen, Animal Models for Influenza Virus Pathogenesis and Transmission. Viruses 2, 1530–1563 (2010).

15. E. P. Rock, P. R. Sibbald, M. M. Davis, Y. H. Chien, CDR3 length in antigen-specific immune receptors. J Exp Med 179, 323–328 (1994).

16. K. Wiehe, D. Easterhoff, K. Luo et al., Antibody Light-Chain-Restricted Recognition of the Site of Immune Pressure in the RV144 HIV-1 Vaccine Trial Is Phylogenetically Conserved. Immunity 41, 909–918 (2014).

17. D. M. Dudley, M. T. Aliota, E. L. Mohr et al., A rhesus macaque model of Asian-lineage Zika virus infection. Nature communications 7, 12204 (2016).

18. G. W. Dick, S. F. Kitchen, A. J. Haddow, Zika virus. I. Isolations and serological specificity. Trans R Soc Trop Med Hyg 46, 509–520 (1952).

19. K. M. Bialas, T. Tanaka, D. Tran et al., Maternal CD4+ T cells protect against severe congenital cytomegalovirus disease in a novel nonhuman primate model of placental cytomegalovirus transmission. Proc Natl Acad Sci U S A 112, 13645–13650 (2015).

20. R. Zhang, L. Verkoczy, K. Wiehe et al., Initiation of immune tolerance-controlled HIV gp41 neutralizing B cell lineages. Sci Transl Med 8, 336ra362 (2016).

21. R. W. Sanders, M. J. van Gils, R. Derking et al., HIV-1 VACCINES. HIV-1 neutralizing antibodies induced by native-like envelope trimers. Science 349, aac4223 (2015).

22. D. Corti, A. L. Suguitan, Jr., D. Pinna et al., Heterosubtypic neutralizing antibodies are produced by individuals immunized with a seasonal influenza vaccine. J Clin Invest 120, 1663–1673 (2010).

23. M. Throsby, E. van den Brink, M. Jongeneelen et al., Heterosubtypic neutralizing monoclonal antibodies cross-protective against H5N1 and H1N1 recovered from human IgM+ memory B cells. PLoS One 3, e3942 (2008).

24. M. A. Moody, N. L. Yates, J. D. Amos et al., HIV-1 gp120 vaccine induces affinity maturation in both new and persistent antibody clonal lineages. J Virol 86, 7496–7507 (2012).

25. E. S. Gray, M. A. Moody, C. K. Wibmer et al., Isolation of a monoclonal antibody that targets the alpha-2 helix of gp120 and represents the initial autologous neutralizing-antibody response in an HIV-1 subtype C-infected individual. J Virol 85, 7719–7729 (2011).

26. M. A. Moody, R. Zhang, E. B. Walter et al., H3N2 influenza infection elicits more cross-reactive and less clonally expanded anti-hemagglutinin antibodies than influenza vaccination. PLoS One 6, e25797 (2011).

27. A. Punjani, H. Zhang, D. J. Fleet, Non-uniform refinement: adaptive regularization improves single-particle cryo-EM reconstruction. Nat Methods 17, 1214–1221 (2020).

28. A. Punjani, J. L. Rubinstein, D. J. Fleet, M. A. Brubaker, cryoSPARC: algorithms for rapid unsupervised cryo-EM structure determination. Nat Methods 14, 290–296 (2017).

29. M. G. Joyce, A. K. Wheatley, P. V. Thomas et al., Vaccine-Induced Antibodies that Neutralize Group 1 and Group 2 Influenza A Viruses. Cell 166, 609–623 (2016).

30. N. L. Kallewaard, D. Corti, P. J. Collins et al., Structure and Function Analysis of an Antibody Recognizing All Influenza A Subtypes. Cell 166, 596–608 (2016).

31. A. Watanabe, K. R. McCarthy, M. Kuraoka et al., Antibodies to a Conserved Influenza Head Interface Epitope Protect by an IgG Subtype-Dependent Mechanism. Cell 177, 1124–1135.e1116 (2019).

32. C. J. Wei, J. C. Boyington, K. Dai et al., Cross-neutralization of 1918 and 2009 influenza viruses: role of glycans in viral evolution and vaccine design. Sci Transl Med 2, 24ra21 (2010).

33. D. J. DiLillo, P. Palese, P. C. Wilson, J. V. Ravetch, Broadly neutralizing anti-influenza antibodies require Fc receptor engagement for in vivo protection. J Clin Invest 126, 605–610 (2016).

34. D. J. DiLillo, G. S. Tan, P. Palese, J. V. Ravetch, Broadly neutralizing hemagglutinin stalk-specific antibodies require FcγR interactions for protection against influenza virus in vivo. Nat Med 20, 143–151 (2014).

35. D. Corti, J. Voss, S. J. Gamblin et al., A neutralizing antibody selected from plasma cells that binds to group 1 and group 2 influenza A hemagglutinins. Science 333, 850–856 (2011).

36. T. Bizebard, B. Gigant, P. Rigolet et al., Structure of influenza virus haemagglutinin complexed with a neutralizing antibody. Nature 376, 92–94 (1995).

37. L. A. Grohskopf, J. M. Ferdinands, L. H. Blanton, K. R. Broder, J. Loehr, Prevention and Control of Seasonal Influenza with Vaccines: Recommendations of the Advisory Committee on Immunization Practices - United States, 2024-25 Influenza Season. MMWR Recomm Rep 73, 1–25 (2024).

38. M. T. Osterholm, N. S. Kelley, A. Sommer, E. A. Belongia, Efficacy and effectiveness of influenza vaccines: a systematic review and meta-analysis. Lancet Infect Dis 12, 36–44 (2012).

39. F. Krammer, G. J. D. Smith, R. A. M. Fouchier et al., Influenza. Nat Rev Dis Primers 4, 3 (2018).

40. T. R. Mosmann, A. J. McMichael, A. LeVert, J. W. McCauley, J. W. Almond, Opportunities and challenges for T cell-based influenza vaccines. Nature reviews. Immunology, (2024).

41. C. Jiao, B. Wang, P. Chen, Y. Jiang, J. Liu, Analysis of the conserved protective epitopes of hemagglutinin on influenza A viruses. Front Immunol 14, 1086297 (2023).

42. A. Gupta, A. Rudra, K. Reed, R. Langer, D. G. Anderson, Advanced technologies for the development of infectious disease vaccines. Nat Rev Drug Discov, (2024).

43. J. R. Whittle, A. K. Wheatley, L. Wu et al., Flow cytometry reveals that H5N1 vaccination elicits cross-reactive stem-directed antibodies from multiple Ig heavy-chain lineages. J Virol 88, 4047–4057 (2014).

44. B. F. Haynes, M. A. Moody, M. Alam et al., Progress in HIV-1 vaccine development. J Allergy Clin Immunol 134, 3–10; quiz 11 (2014).

45. R. A. Gibbs, J. Rogers, M. G. Katze et al., Evolutionary and biomedical insights from the rhesus macaque genome. Science 316, 222–234 (2007).

46. N. Darricarrère, Y. Qiu, M. Kanekiyo et al., Broad neutralization of H1 and H3 viruses by adjuvanted influenza HA stem vaccines in nonhuman primates. Sci Transl Med 13, (2021).

47. H. X. Tan, S. Jegaskanda, J. A. Juno et al., Subdominance and poor intrinsic immunogenicity limit humoral immunity targeting influenza HA stem. J Clin Invest 129, 850–862 (2019).

48. W. C. Liu, J. T. Jan, Y. J. Huang, T. H. Chen, S. C. Wu, Unmasking Stem-Specific Neutralizing Epitopes by Abolishing N-Linked Glycosylation Sites of Influenza Virus Hemagglutinin Proteins for Vaccine Design. J Virol 90, 8496–8508 (2016).

49. A. K. Harris, J. R. Meyerson, Y. Matsuoka et al., Structure and accessibility of HA trimers on intact 2009 H1N1 pandemic influenza virus to stem region-specific neutralizing antibodies. Proc Natl Acad Sci U S A 110, 4592–4597 (2013).

50. T. M. Uyeki, S. Milton, C. Abdul Hamid et al., Highly Pathogenic Avian Influenza A(H5N1) Virus Infection in a Dairy Farm Worker. N Engl J Med 390, 2028–2029 (2024).

## Reference

1. G. Ott, G. L. Barchfeld, G. Van Nest, Enhancement of humoral response against human influenza vaccine with the simple submicron oil/water emulsion adjuvant MF59. Vaccine 13, 1557–1562 (1995).

2. D. C. Ekiert et al., Antibody recognition of a highly conserved influenza virus epitope. Science 324, 246–251 (2009).

3. M. Throsby et al., Heterosubtypic neutralizing monoclonal antibodies cross-protective against H5N1 and H1N1 recovered from human IgM+ memory B cells. PLoS One 3, e3942 (2008).

4. J. R. Whittle et al., Broadly neutralizing human antibody that recognizes the receptor-binding pocket of influenza virus hemagglutinin. Proc Natl Acad Sci U S A 108, 14216–14221 (2011).

5. M. A. Moody et al., H3N2 influenza infection elicits more cross-reactive and less clonally expanded anti-hemagglutinin antibodies than influenza vaccination. PLoS One 6, e25797 (2011).

6. A. K. Harris et al., Structure and accessibility of HA trimers on intact 2009 H1N1 pandemic influenza virus to stem region-specific neutralizing antibodies. Proc Natl Acad Sci U S A 110, 4592–4597 (2013).

7. S. Johnson et al., Development of a humanized monoclonal antibody (MEDI-493) with potent in vitro and in vivo activity against respiratory syncytial virus. J Infect Dis 176, 1215–1224 (1997).

8. S. M. Alam et al., Human immunodeficiency virus type 1 gp41 antibodies that mask membrane proximal region epitopes: antibody binding kinetics, induction, and potential for regulation in acute infection. J Virol 82, 115–125 (2008).

9. H. X. Liao et al., High-throughput isolation of immunoglobulin genes from single human B cells and expression as monoclonal antibodies. J Virol Methods 158, 171–179 (2009).

10. H. X. Liao et al., Vaccine induction of antibodies against a structurally heterogeneous site of immune pressure within HIV-1 envelope protein variable regions 1 and 2. Immunity 38, 176–186 (2013).

11. E. S. Gray et al., Antibody specificities associated with neutralization breadth in plasma from human immunodeficiency virus type 1 subtype C-infected blood donors. J Virol 83, 8925–8937 (2009).

12. R. Cottey, C. A. Rowe, B. S. Bender, Influenza virus. Curr Protoc Immunol Chapter 19, Unit 19.11 (2001).

13. M. A. Moody et al., HIV-1 gp120 vaccine induces affinity maturation in both new and persistent antibody clonal lineages. J Virol 86, 7496–7507 (2012).

14. K. Wiehe et al., Antibody light-chain-restricted recognition of the site of immune pressure in the RV144 HIV-1 vaccine trial is phylogenetically conserved. Immunity 41, 909–918 (2014).

15. J. M. Volpe, L. G. Cowell, T. B. Kepler, SoDA: implementation of a 3D alignment algorithm for inference of antigen receptor recombinations. Bioinformatics 22, 438–444 (2006).

16. T. B. Kepler et al., Reconstructing a B-Cell Clonal Lineage. II. Mutation, Selection, and Affinity Maturation. Front Immunol 5, 170 (2014).

17. T. B. Kepler, Reconstructing a B-cell clonal lineage. I. Statistical inference of unobserved ancestors. F1000Res 2, 103 (2013).

18. C. J. Wei et al., Cross-neutralization of 1918 and 2009 influenza viruses: role of glycans in viral evolution and vaccine design. Sci Transl Med 2, 24ra21 (2010).

19. H. Xu et al., Key mutations stabilize antigen-binding conformation during affinity maturation of a broadly neutralizing influenza antibody lineage. Proteins 83, 771–780 (2015).

20. A. G. Schmidt et al., Preconfiguration of the antigen-binding site during affinity maturation of a broadly neutralizing influenza virus antibody. Proc Natl Acad Sci U S A 110, 264–269 (2013).

21. D. D. Raymond et al., Influenza immunization elicits antibodies specific for an egg-adapted vaccine strain. Nat Med 22, 1465–1469 (2016).

22. D. N. Mastronarde, Automated electron microscope tomography using robust prediction of specimen movements. J Struct Biol 152, 36–51 (2005).

23. A. Punjani, J. L. Rubinstein, D. J. Fleet, M. A. Brubaker, cryoSPARC: algorithms for rapid unsupervised cryo-EM structure determination. Nat Methods 14, 290–296 (2017).

24. A. Punjani, H. Zhang, D. J. Fleet, Non-uniform refinement: adaptive regularization improves single-particle cryo-EM reconstruction. Nat Methods 17, 1214–1221 (2020).

25. T. D. Goddard et al., UCSF ChimeraX: Meeting modern challenges in visualization and analysis. Protein Sci 27, 14–25 (2018).

26. E. F. Pettersen et al., UCSF ChimeraX: Structure visualization for researchers, educators, and developers. Protein Sci 30, 70–82 (2021).

27. S. M. Moin et al., Co-immunization with hemagglutinin stem immunogens elicits cross-group neutralizing antibodies and broad protection against influenza A viruses. Immunity 55, 2405–2418.e2407 (2022).

28. P. Emsley, B. Lohkamp, W. G. Scott, K. Cowtan, Features and development of Coot. Acta Crystallogr D Biol Crystallogr 66, 486–501 (2010).

29. P. D. Adams et al., PHENIX: a comprehensive Python-based system for macromolecular structure solution. Acta Crystallogr D Biol Crystallogr 66, 213–221 (2010).

30. P. V. Afonine et al., Real-space refinement in PHENIX for cryo-EM and crystallography. Acta Crystallogr D Struct Biol 74, 531–544 (2018).

31. K. R. Abhinandan, A. C. Martin, Analysis and improvements to Kabat and structurally correct numbering of antibody variable domains. Mol Immunol 45, 3832–3839 (2008).

32. A. Morin et al., Collaboration gets the most out of software. Elife 2, e01456 (2013).

