## Supplementary figures and images for "Elicitation of stem-directed antibodies in rhesus macaques by a conventional hemagglutinin immunogen"

### Supplemental Figures

#### Supplemental Figure 1

A

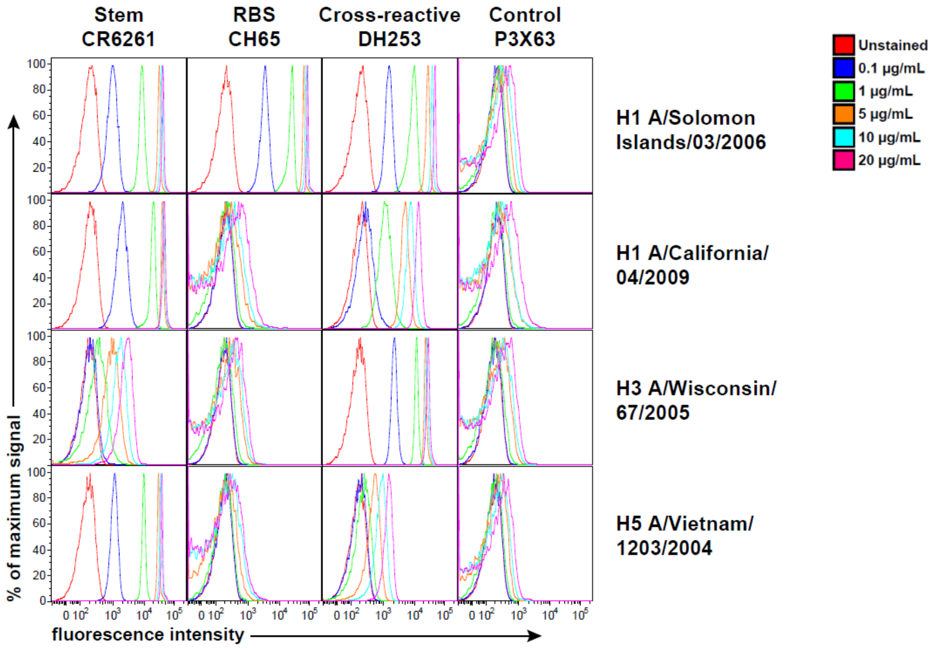


B

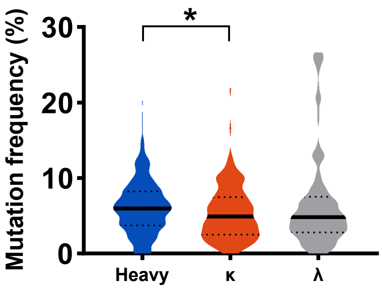

**Supplemental Figure 2**

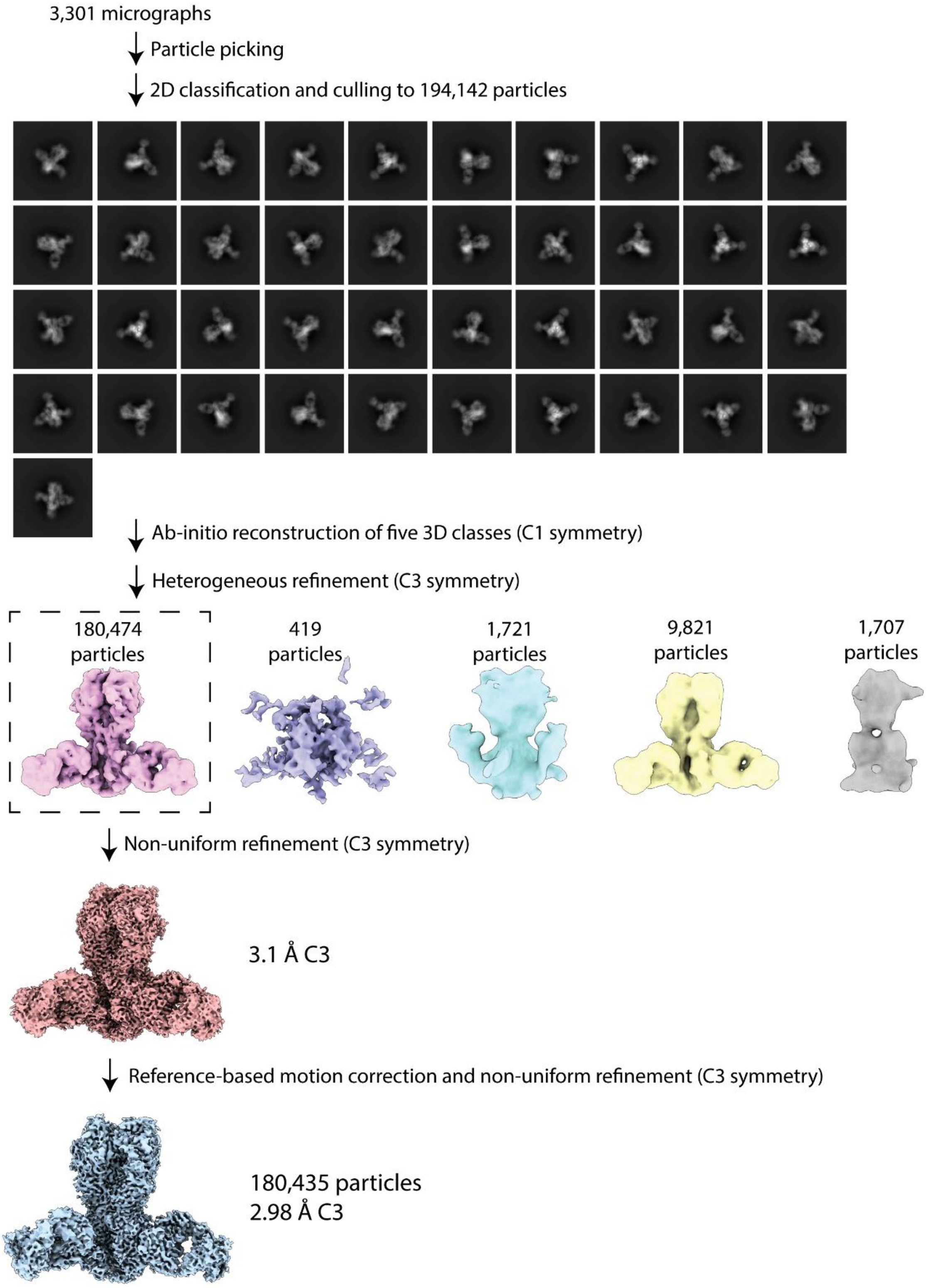

**Supplemental Figure 3**

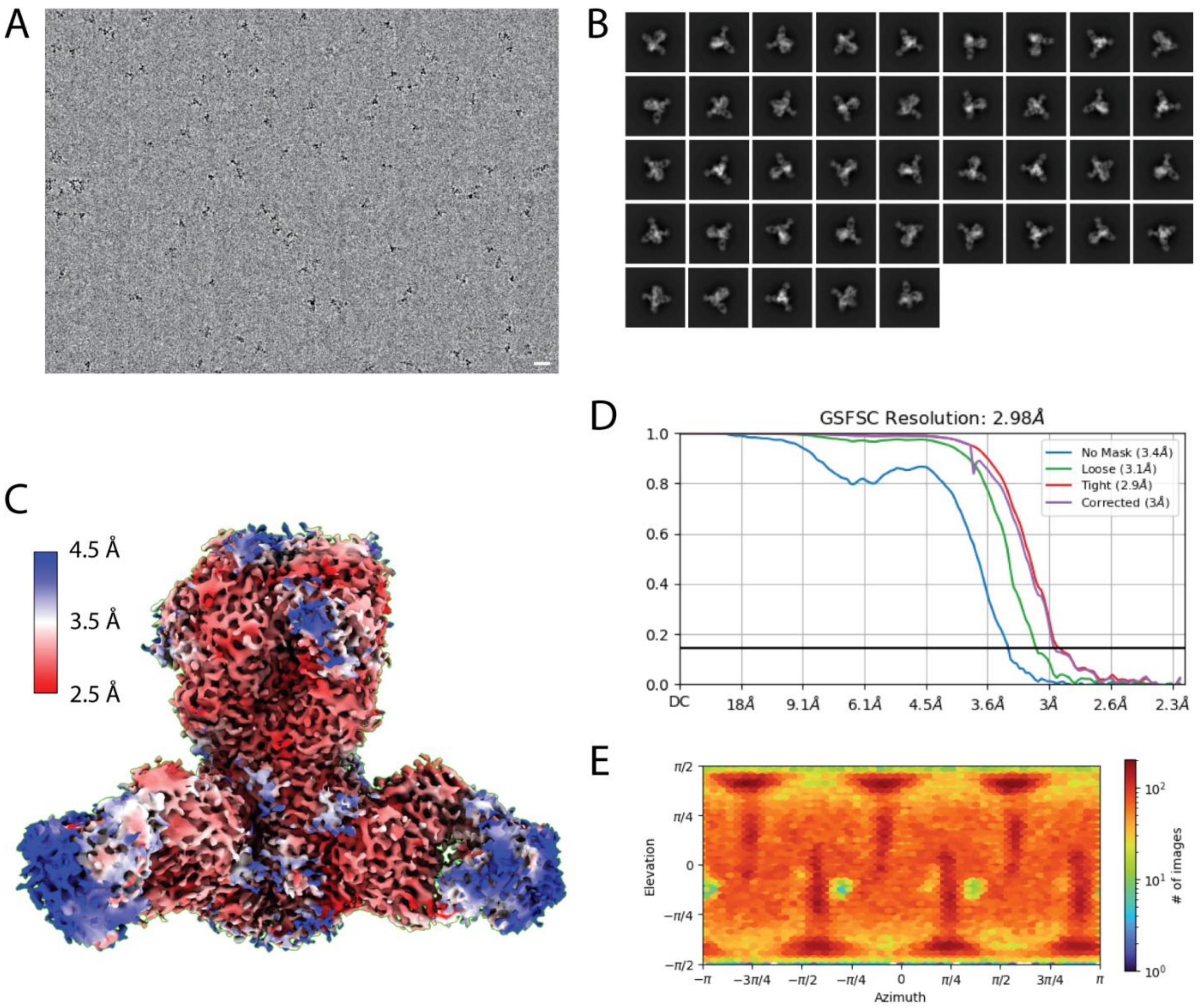

#### Supplemental Figure 4

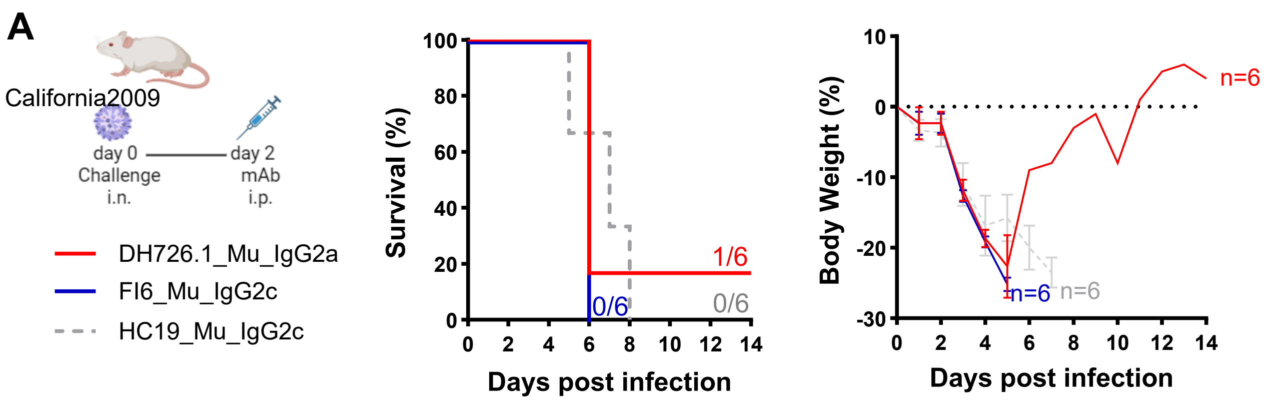


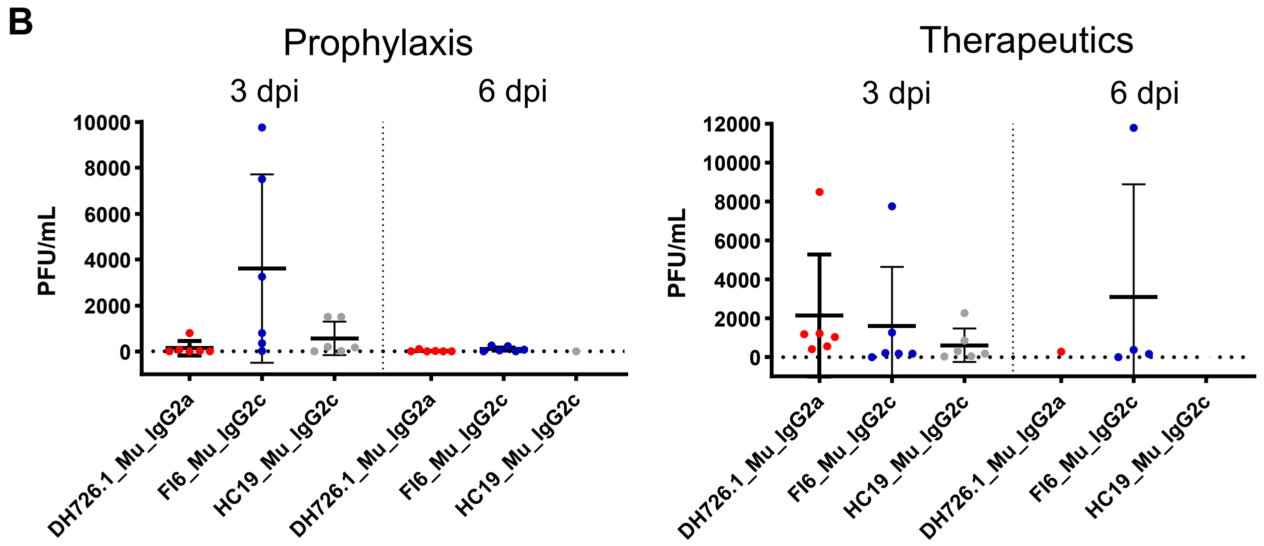


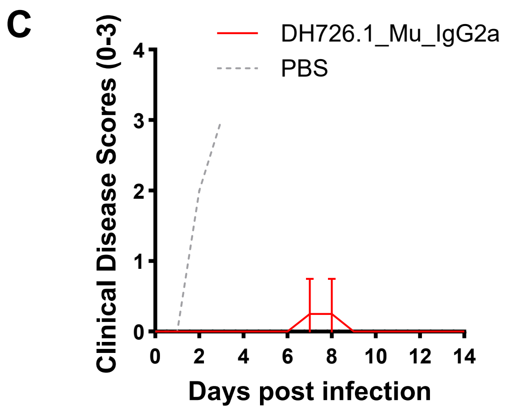
