## Supplemental Tables for "Elicitation of stem-directed antibodies in rhesus macaques by a conventional hemagglutinin immunogen"

**Supplemental Table 1: Cryo-EM data collection and processing statistics**

| Data collection and processing | |
|---|---|
| Magnification (x) | 36000 |
| Voltage (kV) | 200 |
| Electron exposure ($e^-/Å^{-2}$) | 49.99 |
| Defocus range (μm) | 0.8, 2.5 |
| Pixel size (Å) | 1.1 |
| Symmetry imposed | C3 |
| Initial particle images (no.) | 299430 |
| Final particle images (no.) | 180435 |
| Map Resolution (Å) | 2.98 |
| FSC threshold | 0.143 |
| Map resolution range (Å) | 2.5 – 4.5 |

**Supplemental Table 2: H-bond donor/acceptor atoms and distances in the atomic model of DH726.1 Fab bound to H1/SI06**

| Hydrogen bond | H1/SI06 | Bond distance (Å) | DH726.1 Fab HC |
|---|---|---|---|
| 1 | Tyr363 N | 3.1 | Gly53 O |
| 2 | Asp348 OD2 | 3.3 | Asn52 ND2 |
| 3 | Tyr363 O | 3.0 | Asn52 ND2 |
| 4 | Tyr363 O | 3.1 | Gly55 N |
| 5 | Lys482 NZ | 3.4 | Gly55 O |
| 6 | Val347 O | 3.1 | Val100D N |
